# MIAAIM: Multi-omics image integration and tissue state mapping using topological data analysis and cobordism learning

**DOI:** 10.1101/2021.12.20.472858

**Authors:** Joshua M. Hess, Iulian Ilieş, Denis Schapiro, John J. Iskra, Walid M. Abdelmoula, Michael S. Regan, Georgios Theocharidis, Chin Lee Wu, Aristidis Veves, Nathalie Y.R. Agar, Ann E. Sluder, Mark C. Poznansky, Patrick M. Reeves, Ruxandra F. Sîrbulescu

**Affiliations:** Vaccine and Immunotherapy Center, Massachusetts General Hospital, Harvard Medical School, Boston, MA, USA; Tri-Institutional Training Program in Computational Biology and Medicine, Weill Cornell Medicine, New York, NY, USA; Healthcare Systems Engineering Institute, Northeastern University, Boston, MA, USA; Laboratory of Systems Pharmacology, Harvard Medical School, Boston, MA, USA; Broad Institute of Harvard Medical School and Massachusetts Institute of Technology, Boston, MA, USA; Institute for Computational Biomedicine, Faculty of Medicine, Heidelberg University Hospital and Heidelberg University, Heidelberg, Germany; Institute of Pathology, Heidelberg University Hospital, Heidelberg, Germany; Department of Mathematics, Emory & Henry College, Emory, VA; Department of Neurosurgery, Brigham and Women’s Hospital, Harvard Medical School, Boston, MA, USA; Invicro, LLC, Boston, MA, USA; The Rongxiang Xu Center for Regenerative Therapeutics, Beth Israel Deaconess Medical Center, Harvard Medical School, Boston, MA, USA; Department of Pathology, Massachusetts General Hospital, Harvard Medical School, Boston, MA, USA; Department of Radiology, Brigham and Women’s Hospital, Harvard Medical School, Boston, MA, USA

## Abstract

High-parameter tissue imaging enables detailed molecular analysis of single cells in their spatial environment. However, the comprehensive characterization and mapping of tissue states through multimodal imaging across different physiological and pathological conditions requires data integration across multiple imaging systems. Here, we introduce MIAAIM (Multi-omics Image Alignment and Analysis by Information Manifolds) a modular, reproducible computational framework for aligning data across bioimaging technologies, modeling continuities in tissue states, and translating multimodal measures across tissue types. We demonstrate MIAAIM’s workflows across diverse imaging platforms, including histological stains, imaging mass cytometry, and mass spectrometry imaging, to link cellular phenotypic states with molecular microenvironments in clinical biopsies from multiple tissue types with high cellular complexity. MIAAIM provides a robust foundation for the development of computational methods to integrate multimodal, high-parameter tissue imaging data and enable downstream computational and statistical interrogation of tissue states.

## INTRODUCTION

Increased availability of multiplexed imaging at cellular and subcellular resolution allows multiparameter molecular descriptions of organelle, cell, and tissue organization in a preserved spatial context^1–6^. A variety of high-parameter imaging technologies have been developed, each capturing individual aspects of multi-faceted tissue representations – ranging from untargeted methods such as mass spectrometry imaging^7^ (MSI) and spatial transcriptomics^8, 9^, which provide comprehensive molecular profiles, to targeted, multiplexed antibody-based labeling approaches^1–6^. Compared to advances in data acquisition^10, 11^ and atlas mapping for single-cell multi-omics data in dissociated tissue^12, 13^, analogous computational methods that preserve the spatial structure of tissues remain underdeveloped. Tissue cytoarchitecture and molecular composition are crucial for the comprehensive characterization of biological responses under different physiological conditions or disease states. Most importantly, multi-omics tissue portraits, including single-cell composition and spatial structure, could be used to infer equilibrium and transition states of tissue health and disease.

The integration of data across multiple imaging technologies can present significant challenges, including data set size and high dimensionality. In recent years, methods have been developed to align multimodal imaging data as it becomes increasingly relevant to analyze data acquired across multiple modalities^14, 15^. To date, no approach to transform data from separate imaging modalities into the same coordinate system that generalizes to the growing repertoire of high-parameter imaging systems has been developed. Dimensionality reduction is routinely used to model single-cell data, showing that generating functions of cellular phenotype and gene expression closely follow a nonlinear manifold structure. While manifold-based representations of single-cell systems enable cell state trajectory inference^16–18^, tissue states emerging from cellular behavior are often discretized. Clustering is typically used to partition cells into distinct populations^19, 20^ (i.e., cell types), and tissue states are defined based on overall abundances of cell types. We suggest that nonlinear mapping of a systems-level state space may better enable descriptions of continuous transitions from health to disease. For tissue atlases, such as the Human Tumor Atlas Network^21^ and the HuBMAP Consortium^22^, the transfer of information across tissue states may be used for downstream analysis, as is routinely done in dissociated tissue mapping. Comparable intact tissue mapping calls for a reproducible data integration framework that can be implemented at multiple institutions across computing architectures.

Here, we describe MIAAIM (Multi-omics Image Alignment and Analysis by Information Manifolds), a modular framework consisting of 4 new software packages (see **Methods**, **Key Resources Table**) with reproducible workflows to align, model, and transfer data across high-parameter imaging technologies. MIAAIM contains a manifold-based image registration approach, designed to compress and integrate data with diverse densities and spatial resolutions, that can be generalized to multiple high-parameter bioimaging systems. In addition, we introduce *cobordism learning* as a dimension reduction and domain transfer method that models phenotypic manifolds (i.e., tissue states) as the boundary of a higher-dimensional manifold, called a cobordism^23^, that characterizes continuous transitions between tissue states. We demonstrate the applicability of MIAAIM using multimodal imaging data derived from a combination of mass spectrometry-based imaging technologies, including imaging mass cytometry (IMC) and mass spectrometry imaging (MSI), in addition to classical histology. Using this method, we investigate diverse cellular and molecular landscapes in tissue samples with high cellular complexity, including human tonsillar, diabetic foot ulcer (DFU) and prostate cancer biopsies.

## RESULTS

### MIAAIM workflow

MIAAIM is a sequential workflow aimed at providing comprehensive portraits of tissue states. It includes 4 processing stages: (i) image preprocessing and compression with the high-dimensional image preparation (HDIprep) workflow, (ii) image registration with the high-dimensional image registration (HDIreg) workflow, (iii) tissue state modeling with cobordism approximation and projection (PatchMAP), and (iv) cross-modality information transfer with i-PatchMAP (**Fig. 1**). Image integration in MIAAIM begins with two or more assembled images (level 2 data)^24, 25^ or spatially resolved data sets (**Fig. 1,** assembled images). The size and standardized format of assembled images vary by technology. For example, cyclic fluorescence-based methods (e.g., CODEX^3^, CyCIF^5, 6^) assemble BioFormats/OME-compatible^26^ 20-60-plex full-tissue images after correcting uneven illumination (e.g., BaSiC^27^) and stitching tiles (e.g., ASHLAR^28^); other methods acquire 20 to 100-plex data within specified regions of interest (ROIs) (e.g. MIBI^1^, IMC^2^) in the OME format. Additional methods quantify thousands of parameters at rasterized locations on full tissues or ROIs and are not stored in BioFormats/OME-compatible formats (e.g., the imzML format^29^ often stores MSI data). MIAAIM accepts a variety of input data formats (**Fig. 1**), and its modular design will enable it to incorporate additional formats tailored to new imaging modalities.

**Fig. 1.**
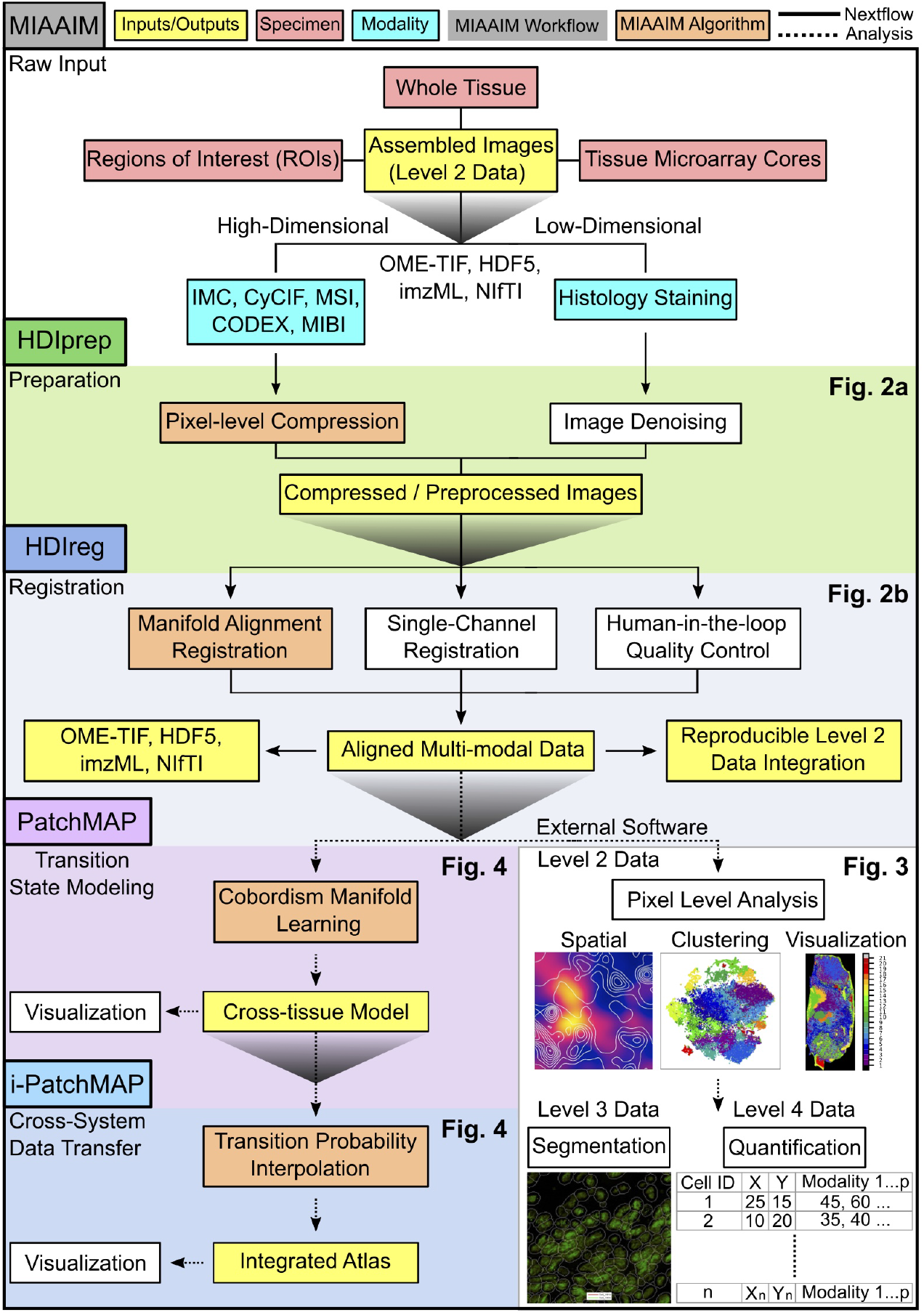
Summary of MIAAIM workflow and key outputs. A schematic representation of the MIAAIM workflow to integrate level 2 multi-modal images derived from various imaging methodologies (cyan boxes) and tissue specimen types (red boxes) to enable tissue state mapping across modalities and tissues. Key inputs and outputs (yellow boxes) of each MIAAIM module (shaded boxes) are connected through MIAAIM’s Nextflow implementation (solid arrows) or exploratory analysis modules in Python (dashed arrows). Algorithms unique to MIAAIM (orange boxes) are detailed in corresponding figures in the remainder of this study (black bolded text). Dataflows associated with high-parameter images (i.e., greater than 3 channels) follow the left-side arrows through the HDIprep (green) and HDIreg (navy blue) modules for pixel-level image compression and pixel manifold-based image registration, while low-parameter images follow the right-side arrows (HDIprep and HDIreg modules, white boxes). Aligned multi-modal data may be exported in various file formats (HDIreg, left-side yellow boxes), and MIAAIM’s Nextflow implementation of the Docker containerized HDIprep and HDIreg modules enables level 2 data integration reproducibility (HDIreg, right-side yellow boxes). MIAAIM’s nonparametric data integration allows for human-in-the-loop quality control of aligned multi-modal data (HDIreg, right-side white box). External analysis techniques that interface with MIAAIM’s key outputs are included in the schematic (bottom right white boxes). Aligned multi-modal data or multi-modal single-cell data (i.e., from cell segmentation) across tissue types and tissue states can be modeled using the PatchMAP module (pink), and data can be transferred across tissue states and modalities using the i-PatchMAP module (light blue).

Regardless of technology, assembled images contain high numbers of heterogeneously distributed parameters, which precludes comprehensive, manually-guided image alignment. In addition, high-dimensional imaging produces large feature spaces that challenge methods used in unsupervised settings. The HDIprep workflow in MIAAIM generates compressed images that preserve multiplex salient features to enable cross-technology statistical comparison while minimizing computational complexity (**Fig. 1,** HDIprep). For images acquired from histological staining, HDIprep provides parallelized smoothing and morphological operations that can be applied sequentially for preprocessing. Image registration with HDIreg produces transformations to combine modalities within the same spatial domain (**Fig. 1**, HDIreg). HDIreg uses Elastix^30^, a parallelized image registration library to calculate transformations, and is optimized to transform large multichannel images with minimal memory use, while also supporting histological stains. HDIreg automates image resizing, padding, and trimming of borders prior to applying image transformations.

Aligned data are well-suited for established single-cell and spatial neighborhood analyses^31^ and they can be segmented to capture multimodal single-cell measures (level 3 and 4 data)^24, 25^, such as average protein expression or spatial features of cells, or analyzed at the pixel level. A common goal in pathology is to use composite tissue portraits to map healthy-to-diseased transitions. To address this, similarities between systems-level tissue states can be visualized with the PatchMAP workflow (**Fig. 1,** PatchMAP). PatchMAP models tissue states as phenotypic manifolds that are stitched to form a higher-dimensional cobordism. The result is a nested model capturing nonlinear intra-system states and cross-system continuities. The cobordism paradigm can be applied as a tissue-based atlas-mapping tool to transfer information across modalities with i-PatchMAP (**Fig. 1,** i-PatchMAP).

MIAAIM’s algorithms are nonparametric, using probability distributions supported by manifolds rather than training data models. MIAAIM is therefore technology-agnostic and generalizes to multiple imaging systems (**Supplementary Table 1**). Nonparametric image registration, however, is an iterative, parameter-tuning process rather than a “black-box” solution. This creates a substantial challenge for reproducible data integration across institutions and computing architectures. We therefore Docker^32^ containerized MIAAIM’s data integration workflows and developed a Nextflow^33^ implementation to document human-in-the-loop processing and remove language-specific dependencies in accordance with the FAIR (finable, accessible, interoperable, and reusable) data stewardship principles^34^. In the following sections, we describe algorithms unique to MIAAIM and their application to multimodal imaging data sets across technologies and diverse tissue specimens.

### High-Dimensional Image Compression with HDIprep

To compress high-parameter images, HDIprep performs dimensionality reduction on pixels using Uniform Manifold Approximation and Projection (UMAP)^35^ (**Fig. 2a**). We performed a stringent comparison of dimensionality reduction algorithms using imaging data sets of human DFU, prostate cancer, and tonsil tissue biopsies acquired using MSI, IMC, and hematoxylin and eosin (H&E). Based on dimensionality reduction benchmarks, UMAP consistently outperformed competing algorithms in its robustness to noise and ability to efficiently preserve data complexity while capturing morphological structure (**Supplementary Figs. 1, 2, 3**).

**Fig. 2.**
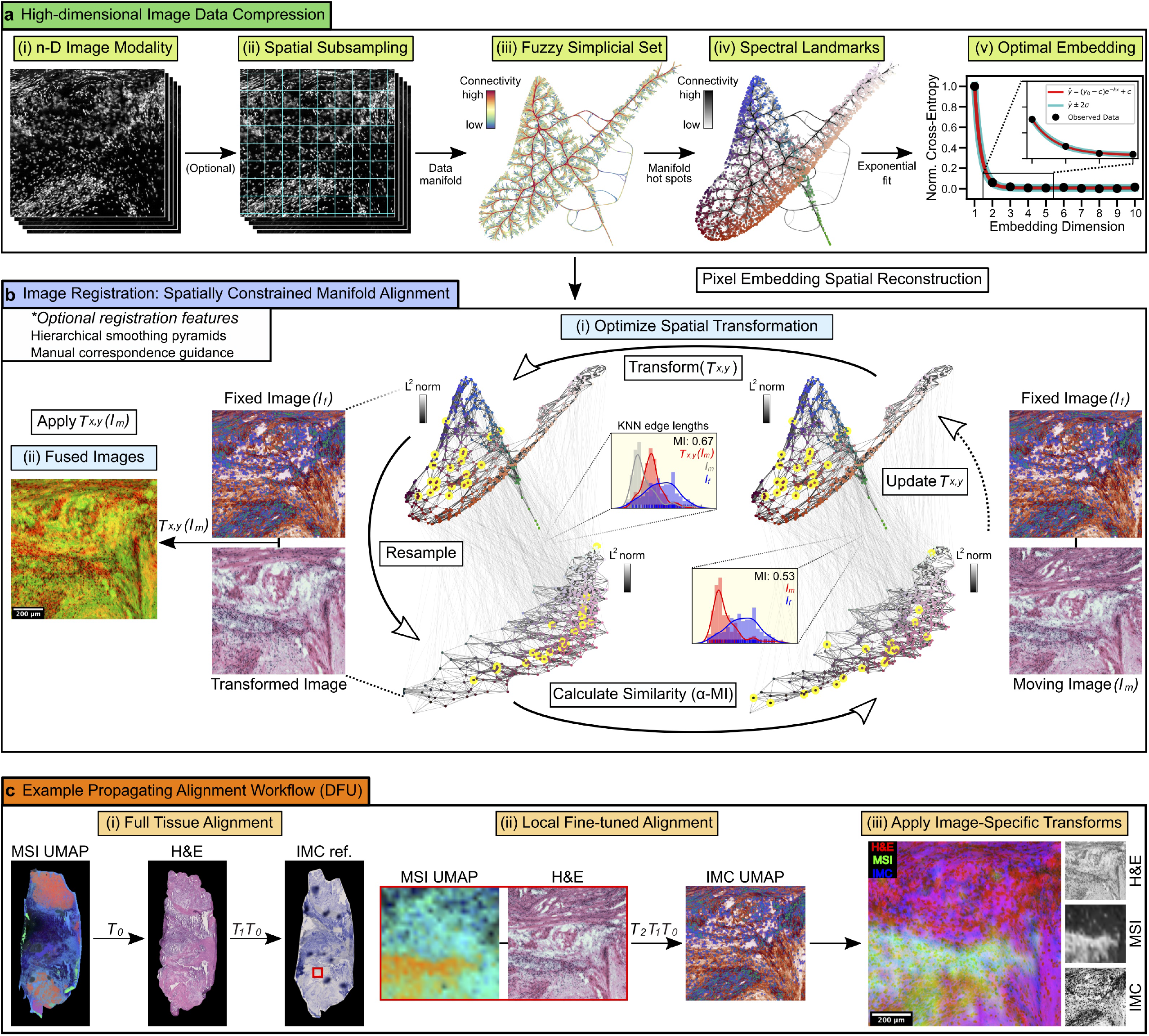
HDIprep compression and spatially constrained manifold alignment with HDIreg. **a.** HDIprep compression algorithm. **(i)** High-dimensional modality. **(ii)** Spatial subsampling. **(iii)** Data manifold represented as a fuzzy simplicial set (UMAP). Edge bundled connectivity of the manifold is shown on two axes of the resulting steady state embedding (*fractal-like structure may not reflect biologically relevant features). **(iv)** High-connectivity landmarks identified with spectral clustering. **(v)** Spectral landmarks are embedded into a range of dimensionalities and exponential regression identifies steady-state dimensionalities. Pixel locations are used to reconstruct compressed image. **b.** HDIreg manifold alignment. **(i)** Spatial transformation is optimized to align moving image to fixed image. KNN graph lengths between resampled points (yellow) are used to compute *α*-MI. Edge-length distribution panels show Shannon MI between distributions of intra-graph edge lengths at resampled locations before and after alignment (*α*-MI converges to Shannon MI as *α* ⟶ 1). MI values show increase in information shared between images after alignment. KNN graph connections show correspondences across modalities. **(ii)** Optimized transformation aligns images. Shown are results of transformed H&E image (green) to IMC (red). **c.** Example alignment. **(i)** Full-tissue MSI-to-H&E registration produces *T*_0_. **(ii)** H&E is transformed to IMC full-tissue reference, producing *T*_1_. **(iii)** ROI coordinates extract underlying MSI and IMC data in IMC reference space. (**iv**) H&E ROI is transformed to correct in IMC domain, producing *T*_2_. Final alignment applies modality-specific transformations. Shown are results for an IMC ROI.

HDIprep retains data complexity with the fewest degrees of freedom necessary by detecting *steady-state manifold embeddings.* To identify steady-state dimensionalities, information captured by UMAP pixel embeddings is computed (**Methods**, **Definition 1,** cross-entropy) across a range of embedding dimensionalities, and the first dimension where the observed cross-entropy approaches the asymptote of an exponential regression fit is selected. Steady state embedding calculations scale quadratically with the number of pixels, HDIprep therefore embeds spectral landmarks in the pixel manifold representative of its global structure (**Supplementary Fig. 4)**.

Pixel-level dimensionality reduction is computationally expensive for large images, i.e., at high resolution (e.g., ≤ 1 *μ*m/pixel). To reduce compression time while preserving quality, we developed a subsampling approach to embed a spatially representative subset of pixels prior to spectral landmark selection and project out-of-sample pixels into embeddings (**Supplementary Figs. 5, 6, 7)**. HDIprep combines all optimizations with a neural-network UMAP implementation^36^ to scale to whole-tissue images. We demonstrate its efficacy on publicly available^37^ 44-channel CyCIF images containing ~100 and ~256 million pixels (**Supplementary Fig. 8).** Thus, HDIprep presents an objective, pixel-level compression method applicable to multiple modalities (**Methods, Algorithm 1**).

### High-Dimensional Image Registration (HDIreg)

MIAAIM connects the HDIprep and HDIreg workflows with a manifold alignment scheme parametrized by spatial transformations. To align pixel manifolds, we extended methodology to compute manifold *α*-entropy^38, 39^ using entropic graphs to UMAP embeddings and adapted it to image registration using the entropic graph-based Rényi *α*-mutual information (*α*-MI)^40^ (**Methods, HDIreg**). HDIreg selects for a transformation that maximizes image-to-image (manifold-to-manifold) *α*-MI (**Fig. 2b**). The entropic-graph based *α*-MI generalizes to arbitrary dimensionalities by considering distributions of k-nearest neighbor (KNN) graph lengths of compressed (i.e., embedded) pixels, rather than directly comparing the pixels themselves. Combining HDIprep compression with KNN *α*-MI extends intensity-based registration to images across technologies, without the need to match specific markers or analytes.

### MIAAIM generates information on cellular phenotype, molecular ion distribution, and tissue state across scales

To demonstrate the capacity of MIAAIM to generate linked data between multimodal imaging with various scales and data densities, we applied the HDIprep and HDIreg workflows to MALDI-TOF MSI, H&E and IMC data from a DFU tissue biopsy containing a spectrum of tissue states, from the necrotic center of the ulcer to the healthy margin. Image acquisition covered 1.2 cm^2^ for H&E and MSI data. Molecular imaging with MSI enabled untargeted mapping of lipids and small metabolites in the 400-1000 *m/z* range across the specimen at a resolution of 50 *μ*m/pixel. Tissue morphology was captured with H&E at 0.2 *μ*m/pixel, and 27-plex IMC data was acquired at 1 *μ*m/pixel resolution from 7 ROIs on an adjacent section.

Cross-modality alignment was performed in a global-to-local fashion **(Fig. 2c)**. We utilized HDIprep compression for high-parameter data and HDIreg manifold alignment for registration of compressed images. Due to the destructive nature of IMC acquisition^2^, we aligned full-tissue data from MSI, H&E (down-sampled to approximately 3.5 *μ*m/pixel), and an IMC reference image first. Deformations not captured at the full-tissue scale within each ROI were corrected using manual landmark guidance. Tissue deformations resulting from serial sectioning were accounted for with nonlinear transformations. Registrations were initialized by affine transformations for coarse alignment before nonlinear correction. Resolution differences were accounted for with a multiresolution smoothing scheme^30^. Final alignment proceeded by composing modality and ROI-specific transformations.

After locating single-cells using image segmentation with Ilastik^41^, single-cell parameter quantification with MCMICRO^24^, and antibody staining quality control, registered images yielded the following information for 7,114 cells: (i) average expression of 14 proteins (**Methods, Key Resources Table**) including markers for lymphocytes, macrophages, fibroblasts, keratinocytes, and endothelial cells, as well as extracellular matrix proteins, such as collagen and smooth muscle actin; (ii) morphological features, such as cell eccentricity, solidity, extent, and area, spatial positioning of each cell centroid; and (iii) the distribution of 9,753 *m/z* MSI peaks across the full tissue. Distances from each MSI pixel and IMC ROI to the center of the ulcer, identified by manual inspection of H&E, were also quantified. Through the integration of these modalities, MIAAIM provided cross-modal information that could not be gathered with a single imaging system, such as profiling of single-cell protein expression and microenvironmental molecular abundance.

### Identification of molecular microenvironmental niches correlated with cell and disease states

We verified the existence of cross-modal associations from the integrated multimodal imaging data set by conducting a *microenvironmental correlation network analysis* (MCNA) on registered IMC and MSI data (**Fig. 3**). We performed community detection (i.e., clustering) on MSI analytes (*m/z* peaks) based on their correlations to single-cell protein measures and defined *microenvironmental correlation network modules* (MCNMs; different colors in **Fig. 3a**). Inspection of MCNMs with top correlations to protein levels identified with IMC revealed that sets of molecules, rather than individual peaks, were associated with cellular protein expression (**Fig. 3b**). MCNMs organized on an axis separating those with moderate positive correlations to cell markers indicative of inflammation and cell death (CD68, activated Caspase-3) and those with moderate positive correlations to markers of immune regulation (CD163, CD4, FoxP3) and vasculature (CD31). Some proteins, such as the myeloid marker CD14 and the cell proliferation marker Ki-67, were not strongly correlated with any of the detected *m/z* peaks across all cells.

**Fig. 3.**
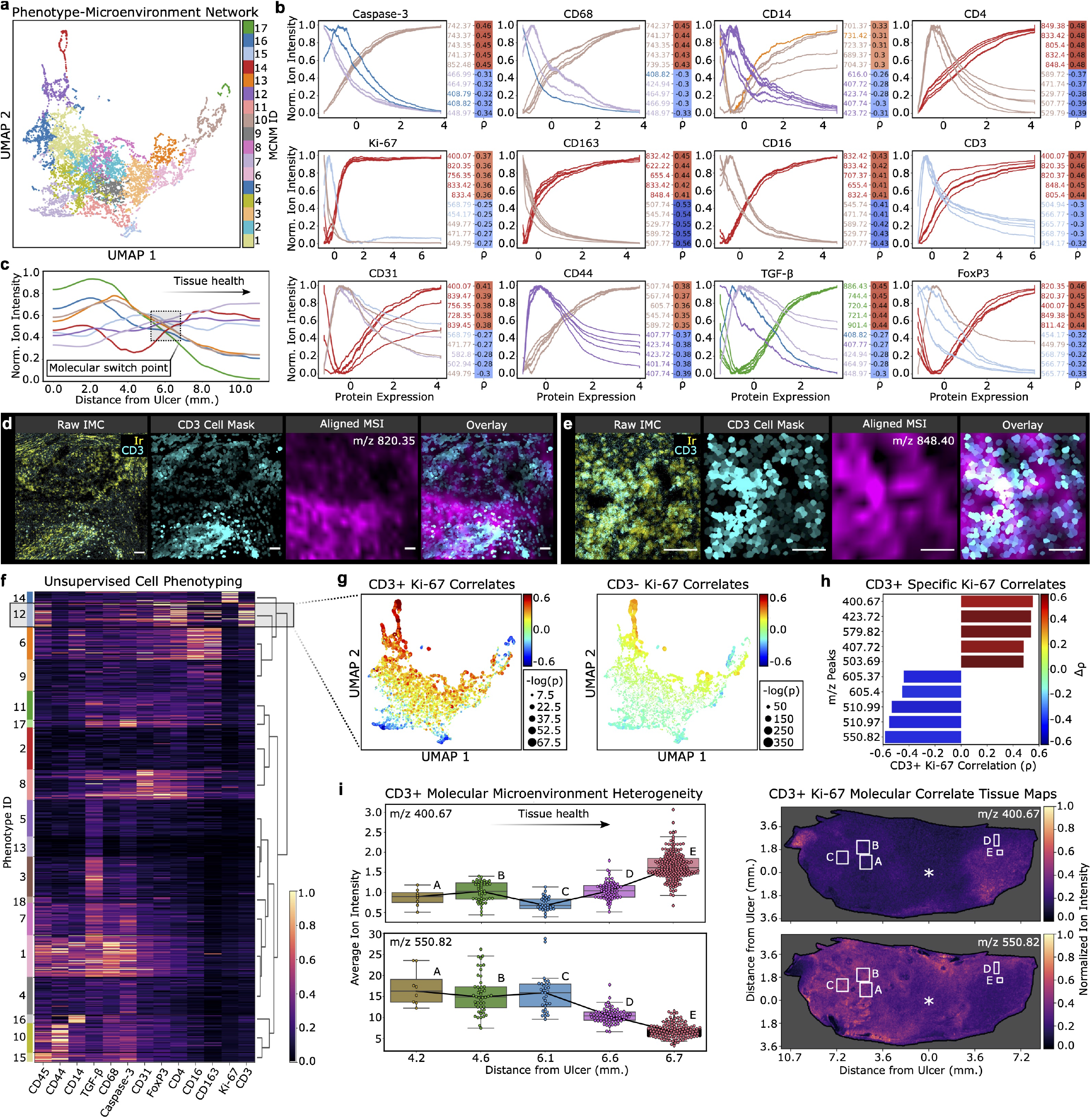
Microenvironmental correlation network analysis (MCNA) links protein expression with molecular distributions in the DFU niche. **a.** MCNA UMAP of *m/z* peaks grouped into modules. **b.** Exponential-weighted moving averages of normalized ion intensities for top five positive and negative correlates to proteins. Colors indicate module assignment. Heatmaps (right) indicate Spearman’s rho. **c.** Exponential-weighted moving averages of normalized average ion intensity per modules ordered as distance from center of wound in DFU increases. **d.** Raw IMC nuclear (Ir) and CD3 staining in ROI (left) (scale bars = 80 *μ*m). Masks showing CD3 expression (middle-left). Aligned MSI showing one of top CD3 correlates (middle-right). Overlay of CD3 expression and a top molecular correlate (right). **e.** Same as **d** at different ROI. **f.** Unsupervised phenotyping. Shaded box indicates CD3+ population. Heatmap indicates normalized protein expression. **g.** MCNA UMAP colored to reflect ions’ correlations to Ki-67 within CD3+ and CD3-populations. Colors indicate Spearman’s rho and size of points indicates negative log transformed, Benjamini-Hochberg corrected P-values for correlations. **h.** Tornado plot showing top five CD3+ differential negative and positive correlates to Ki-67 compared to the CD3 - cell populations. X-axis indicates CD3+ specific Ki-67 values. Color of each bar indicates change in correlation from CD3- to CD3+ populations. **i.** Boxplots showing ion intensity and of top differentially correlated ions (top, positive; bottom; negative) to CD3+ specific Ki-67 expression across ROIs on the DFU. Tissue maps of top differentially associated CD3+ Ki-67 correlates (top, positive; bottom; negative) with boxes (white) indicating ROIs on the tissue that contain CD3+ cells.

To gain insights into the association of molecular distributions with tissue health, we plotted ion intensity distributions of MCNMs with respect to their proximity to the center of the ulcer (**Fig. 3c**). This analysis revealed a change in molecular profiles about 6 mm from the ulcer center point, as the tissue state progressed from healthy to injured. We validated our observations and the performance of HDIreg to align micron-scale structures by visualizing the distribution of top correlated ions within cellular microenvironments (**Figs. 3d, 3e**).

An advantage of the MIAAIM approach is the capacity to identify molecular variations in one modality (here, MSI) that correlate with cell states identified using a different modality (here, IMC). We investigated whether *m/z* peaks differentially associate with cell proliferation (Ki-67 marker in IMC). We identified cell phenotypes through unsupervised clustering on IMC segmented cell-level expression patterns (**Fig. 3f**) and conducted a differential correlation analysis between phenotypes within the well-separated CD3+ cluster, which likely identifies infiltrating T cells at the wound site, and the CD3-cell populations (**Fig. 3g**). Interestingly, we found that correlations to Ki-67 expression shifted with near significance (2*σ*) for multiple *m/z* peaks (Fisher transformed, one-sided z-statistics; Bonferroni corrected P-values) between CD3– populations and the CD3+ population (**Fig. 3h**).

We then utilized spatial context preserved with MIAAIM and observed that ion intensities of *m/z* peaks positively correlated to Ki-67 in CD3+ cells increased with distance from the wound, while molecules with Ki-67 negative correlations specific to CD3+ cells showed the opposite trend (**Fig. 3i**). This distribution of Ki-67 correlates suggests that proliferation of CD3+ T cells occurs predominantly near the healthy margin of the DFU and confirms that molecular correlates of T-cell proliferation can be identified through this unbiased analysis. Collectively, these results provide insights to molecular microenvironments associated with different functional and metabolic states of cell subtypes, and how these microenvironments are distributed in the spatial context in a gradient from injured to healthy tissue.

### Mapping tissue states via cobordism approximation and projection (PatchMAP)

To model transitions between tissue states, such as healthy or injured, we generalized manifold learning to higher-order instances by developing a new algorithm, called PatchMAP, that integrates mutual nearest-neighbor calculations with UMAP **(Fig. 4a** and **Methods, Algorithm 2)**. We hypothesized that state spaces of system-level transitions could be represented as nonlinear manifolds. Accordingly, PatchMAP represents the disjoint union of phenotypic manifolds as the boundary of a nonlinear cobordism that maps the transition between phenotypic manifolds. Patches across boundary manifolds are connected by pairwise nearest-neighbor queries that represent cobordism geodesics and stitched using the t-norm to make their metrics compatible. Interpreting PatchMAP embeddings is analogous to existing dimensionality reduction algorithms – similar data within or across boundary manifolds are located close to each other, while dissimilar data are farther apart. PatchMAP incorporates both boundary manifold topological structure and continuities across boundary manifolds to produce cobordisms.

**Fig. 4.**
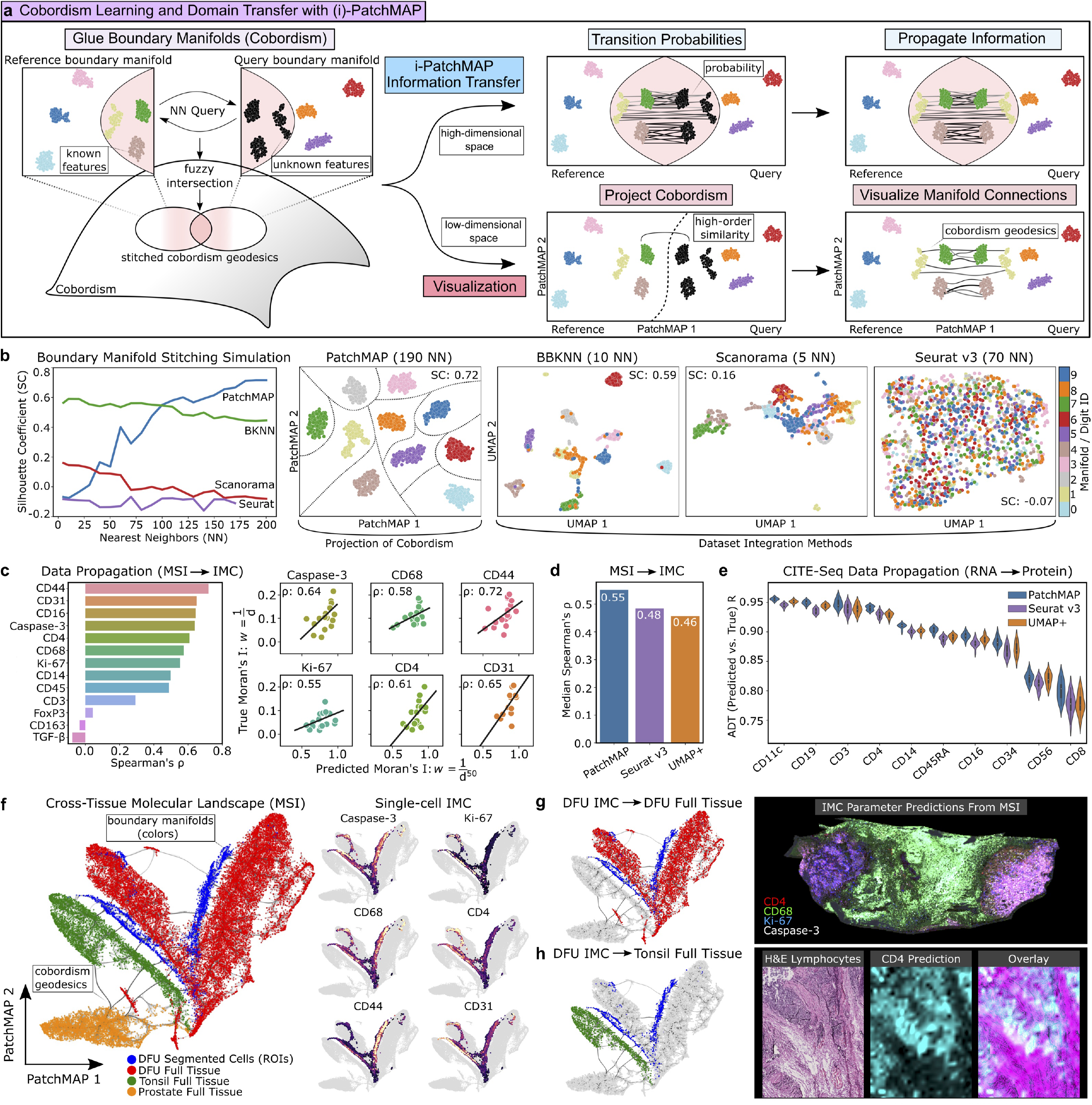
Cobordism projection with PatchMAP and domain transfer with i-PatchMAP. **a.** Schematic representing the PatchMAP algorithm’s cobordism geodesic stitching between boundary manifolds (reference and query data) to form cobordism (represented as grey surface), information transfer across cobordism geodesics with the i-PatchMAP workflow (top right), and cobordism projection visualization (bottom). In PatchMAP, cobordisms sharing common boundary manifolds are glued along their shared domain, and geodesic distances in the cobordism are made compatible with boundary manifold geodesics. **b.** Boundary manifold stitching simulation. PatchMAP projection (manually drawn dashed lines indicate stitching) and UMAP projections of integrated data are shown at NN values that maximized the SC for each method. **c.** MSI-to-IMC data transfer with i-PatchMAP. Line plots show Spearman’s rho between predicted and true spatial autocorrelation values. **d.** MSI-to-IMC data transfer benchmark. **e.** CBMC multimodal CITE-seq data transfer benchmark. **f.** PatchMAP of DFU segmented single-cells (blue), DFU full tissue (red), tonsil tissue (green), and prostate tissue (orange) pixels based on MSI profile. Individual plots show IMC expression for DFU single-cells (right). **g.** MSI-to-IMC data transfer from DFU single-cells to the DFU full tissue using i-PatchMAP. **h.** MSI-to-IMC data transfer from DFU single-cells to the tonsil tissue.

Currently, no machine learning method to form cobordisms exists – methods closest to doing this are single-cell data integration algorithms^13, 42, 43^. Therefore, to benchmark PatchMAP’s manifold stitching, we compared it to the data integration methods BBKNN^43^, Seurat v3^13^, and Scanorama^42^ in a stitching simulation using the “digits” machine learning method development data set (**Fig. 4b**). We split data by label into boundary manifolds, then applied each method to re-stitch the full data set. In this task, perfect stitching would result in the separation of projected boundary manifolds, which we quantified with the silhouette coefficient (SC). For controlled visualization, UMAP was used after data integration to provide analogous embeddings to PatchMAP’s for all benchmark methods.

PatchMAP outperformed data integration methods at higher nearest-neighbor (NN) counts in the stitching simulation (**Fig. 4b**). All other methods incorrectly mixed boundary manifolds when there was no overlap, as expected given that lack of manifold connections violated their assumptions. By contrast, PatchMAP’s stitching uses a fuzzy set intersection^35^, which prunes incorrectly connected data across manifolds while reinforcing correct connections. We also validated that PatchMAP preserves boundary manifold organization while embedding higher-order structures between similar boundary manifolds **(Supplementary Fig. 9)**. At low NN values and when boundary manifolds are similar, PatchMAP resembles UMAP projections (**Supplementary Fig. 9**). At higher NN values, manifold annotations are strongly weighted, which results in less mixing and better manifold separation.

### Information transfer across imaging technologies and tissues (i-PatchMAP)

We posited that the transfer of information across biological states should account for continuous transitions and be robust to manifold connection strength (including the lack thereof). The i-PatchMAP workflow therefore uses PatchMAP as a paired domain transfer and quality control visualization method to propagate information between different imaging methods (**Fig. 4a,** information transfer). To transfer information across data sets, i-PatchMAP first computes one-step Markov chain transition probabilities between boundary manifolds of “*reference*” and “*query*” data along cobordisms (**Fig. 4a,** transition probabilities), and then linearly interpolates measures from reference to query data (**Fig. 4a,** propagate information). Quality control of i-PatchMAP can be performed by visualizing connections between boundary manifolds in PatchMAP embeddings (**Fig. 4a,** visualize manifold connections).

To benchmark i-PatchMAP, we compared it to other nonparametric domain transfer tools, Seurat v3 and a modification of UMAP incorporating a transition probability-based interpolation similar to i-PatchMAP (UMAP+), first on data multimodal single-cell data from DFU ROIs (see **Fig. 3**) and then separately, using a publicly available cord blood mononuclear cell (CBMC) CITE-seq data set^11^. UMAP+ utilized a directed NN graph from query to reference data for data interpolation, rather than PatchMAP’s metric-compatible stitching, therefore acting as a control for PatchMAP. We tiled ROIs from to construct 23 evaluation instances and performed leave-one-out cross-validation using single-cell MSI profiles to predict IMC protein expression. We assessed accuracy using Spearman’s correlation between the predicted spatial autocorrelation (Moran’ s I) and the true autocorrelation for each parameter to account for resolution differences between imaging modalities (**Fig. 4c**). i-PatchMAP outperformed tested methods on its ability to transfer IMC measures to query data based on MSI profiles (**Fig. 4d**). However, all methods demonstrated consistently poor performance for parameters with no original spatial autocorrelation within tiles (TGF-*β*, FoxP3, CD163). For the CITE-seq data set, we created 15 evaluation instances and used single-cell RNA profiles to predict antibody derived tag (ADT) abundance. We quantified performance using Pearson’s correlation between true and predicted ADT values (**Fig. 4e**), finding that i-PatchMAP performed as well or slightly better than other tested methods for all parameters.

### i-PatchMAP transfers multiplexed protein distributions across tissues based on molecular microenvironmental profiles

To assess whether i-PatchMAP can be used to transfer molecular signature information across imaging modalities and further, across different tissue samples, we used single-cell IMC/MSI protein measures (see **Fig. 3**) to extrapolate IMC information from discrete ROIs to the full DFU sample, as well as to distinct prostate tumor and tonsil specimens, based on MSI profiles. A PatchMAP embedding of single cells in DFU ROIs and individual pixels across tissues based on MSI parameters revealed that single-cell molecular microenvironments in the DFU ROIs provided a good representation of the overall DFU molecular profile (**Fig. 4f**). Accordingly, we used i-PatchMAP to transfer DFU single-cell protein measures from the ROIs to the full DFU tissue based on molecular similarities (**Fig. 4g**). i-PatchMAP predicted that the wound area of the DFU tissue would show high expression levels for CD68, a marker of pro-inflammatory macrophages, and activated Caspase-3, a marker of apoptotic cell death. By contrast, the healthy margin of the DFU biopsy was predicted to contain higher levels of CD4, indicating infiltrating T cells, and the cell proliferation marker Ki-67. Interestingly, the PatchMAP visualization revealed that molecular microenvironments corresponding to specific single-cell measures in the DFU (e.g., CD4) were strongly connected with MSI pixels in the tonsil tissue sample (**Fig. 4h**). In the tonsil tissue, which is lymphocyte rich, i-PatchMAP predictions for CD4 matched well with follicular structure, and areas that lacked cell content were accurately predicted to not contain CD4. By contrast, there were no strong connections between the molecular profiles of the DFU biopsy and the prostate cancer sample, which showed poor immune cell infiltration by IMC examination (**Supplementary Fig. 10**). Thus, in the current datasets, strong inter-sample cellular and molecular connections are supported by the presence or absence of common immune cell populations.

## DISCUSSION

We developed MIAAIM, a modular, reproducible computational framework to align, analyze, and transfer information across high-parameter bioimaging modalities. With MIAAIM we have demonstrated that highly detailed and mathematically grounded structures of high-dimensional bioimaging modalities can be linked and propagated across tissues, tissue states, and modalities. As part of MIAAIM, we have developed a manifold learning algorithm based on cobordism theory to translate multiplexed information across tissues. Our combined IMC-MSI approach provided insights into how molecular microenvironmental niches of individual cells may correlate with their states and localization within tissue sections. We showed MIAAIM’s generalizability across diverse healthy and pathological human tissue types and imaging technologies, including those that generate large, whole-tissue datasets.

In its current version, MIAAIM does not incorporate deep learning, which would require large training sets across multiple imaging technologies for generalized application. To our knowledge, no learning models for multiplexed image registration exist, likely due to the large degree of variability in tissue organization across specimens. Similar limitations likely restrict generalized application of domain transfer neural networks for antibody-free labeling of growing repositories of publicly available images. Improvements in deep learning robustness may make its application routine in these settings – a key will be limiting model complexity through appropriate parametrization and regularization. MIAAIM currently provides a first framework to generate standardized training sets for the development of such methods. Its modular implementation will enable versatile workflow incorporation of new method developments such as those from the deep learning community.

With larger sample sizes, our multimodal imaging approach is well positioned to identify novel biomarkers to support diagnosis and prognosis in clinical cases in a disease-agnostic manner. PatchMAP applied to single-cell data may provide a coordinate system on a manifold *between* single-cell network states (i.e., boundary manifolds) that could be used to infer a “single-cell network phenotypic tree” (i.e., cobordism). Computational views of biological systems have proven to be useful^44^; cobordism theory in this context may indicate a deeper relevance of bridging pure mathematics with modern biology. Indeed, connections between cobordisms and vector spaces are the core of a topological field theory^45^ – translating these concepts to biology may provide a formal, algebraic language to map transitions between cellular system states.

In conclusion, MIAAIM provides a conceptual and practical foundation for novel applications in high-dimensional bioimage integration, interrogating composite tissue states, and transferring data across growing numbers of high-parameter imaging modalities. MIAAIM is technology-agnostic and works with any data in BioFormats/OME-compatible, imzML, NIfTI, and HDF5 formats. We envision MIAAIM to have broad applicability, from single-cell proteomic and transcriptomic imaging to untargeted molecular imaging. The harmonization of these data is poised to provide more comprehensive tissue portraits across health and disease states, and transitions between them.

## METHODS

### MIAAIM implementation

MIAAIM workflows are implemented in Python and connected via the Nextflow pipeline language to enable automated results caching and dynamic processing restarts after alteration of workflow parameters, and to streamline parallelized processing of multiple images. MIAAIM is also available as a Python package. Each data integration workflow is containerized to enable reproducible environments and eliminate any language-specific dependencies. MIAAIM’s output interfaces with a number of existing image analysis software tools (see **Supplementary Note 1**, **Combining MIAAIM with existing bioimaging software**). MIAAIM therefore supplements existing tools rather than replaces them.

### High-dimensional image compression and pre-processing (HDIprep)

HDIprep is implemented by specifying sequential processing steps. Options include image compression for high-parameter data, and filtering and morphological operations for single-channel images. Processed images were exported as 32-bit NIfTI-1 images using the NiBabel library in Python. NIfTI-1 was chosen as the default file format for many of MIAAIM’s operations due to its compatibility with Elastix, ImageJ for visualization, and its memory mapping capability in Python.

To compress high-parameter images, HDIprep identifies a steady-state embedding dimensionality for pixel-level data. Compression is initialized with optional, spatially-guided subsampling to reduce data set size. We then implement UMAP to construct a graph representing the data manifold and its underlying topological structure (*FuzzySimplicialSet, **Algorithm 1***). UMAP aims to optimize an embedding of a high-dimensional fuzzy simplicial set (i.e., a weighted, undirected graph) so that the fuzzy set cross-entropy between the embedded simplicial set and the high-dimensional counterpart is minimized, where the fuzzy set cross-entropy is defined as^35^:

#### Definition 1.

Given a reference set *A* and membership functions *u*: *A* → [0,1], *v*: *A* → [0,1], the *fuzzy set cross-entropy C* of (*A*, *u*) and (*A*, *v*) is defined as:

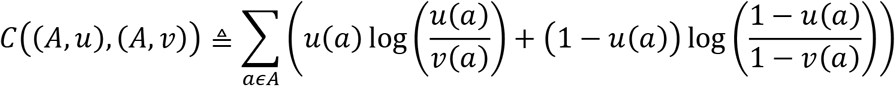

The fuzzy set cross-entropy is a global measure of agreement between simplicial sets, aggregated across members of the reference set *A* (here, graph edges). Calculating its exact value scales quadratically with the number of data points, restricting its use for large data sets. UMAP’s current implementation does not, therefore. compute the exact cross entropy during its optimization of low-dimensional embeddings. Instead, it relies on probabilistic edge sampling and negative sampling to reduce runtimes for large data sets^35^. Congruently, to identify steady-state embedding dimensionalities, we compute patches on the data manifold that are representative of its global structure, and we use these patches in the calculation of the exact cross-entropy after projecting them with UMAP over a range of dimensionalities. The result is a global estimate of the dimensionality required to accurately capture manifold complexity.

To identify globally representative patches on the data manifold, we subject the fuzzy simplicial set to a variant of spectral clustering. We iteratively project the spectral centroids into Euclidean spaces of increasing dimensionality using UMAP and calculate the fuzzy set cross-entropy in each case, then min-max normalize the obtained values. To identify the steady-state embedding dimensionality, we fit a least-squares exponential regression to normalized cross-entropy as a function of dimensionality, then simulate samples along the regression line to find the first dimensionality falling within the 95% confidence interval of the exponential asymptote. The subsampled data is embedded into the steady-state dimensionality, and out-of-sample pixels are projected into this embedding using the native nearest-neighbor based method in UMAP (*transform* () function). Finally, all pixels are mapped back to their original spatial coordinates to construct a compressed image with the number of channels equal to the steady-state embedding dimensionality. These steps are summarized in pseudo-code below:

#### Algorithm 1

Image Compression.

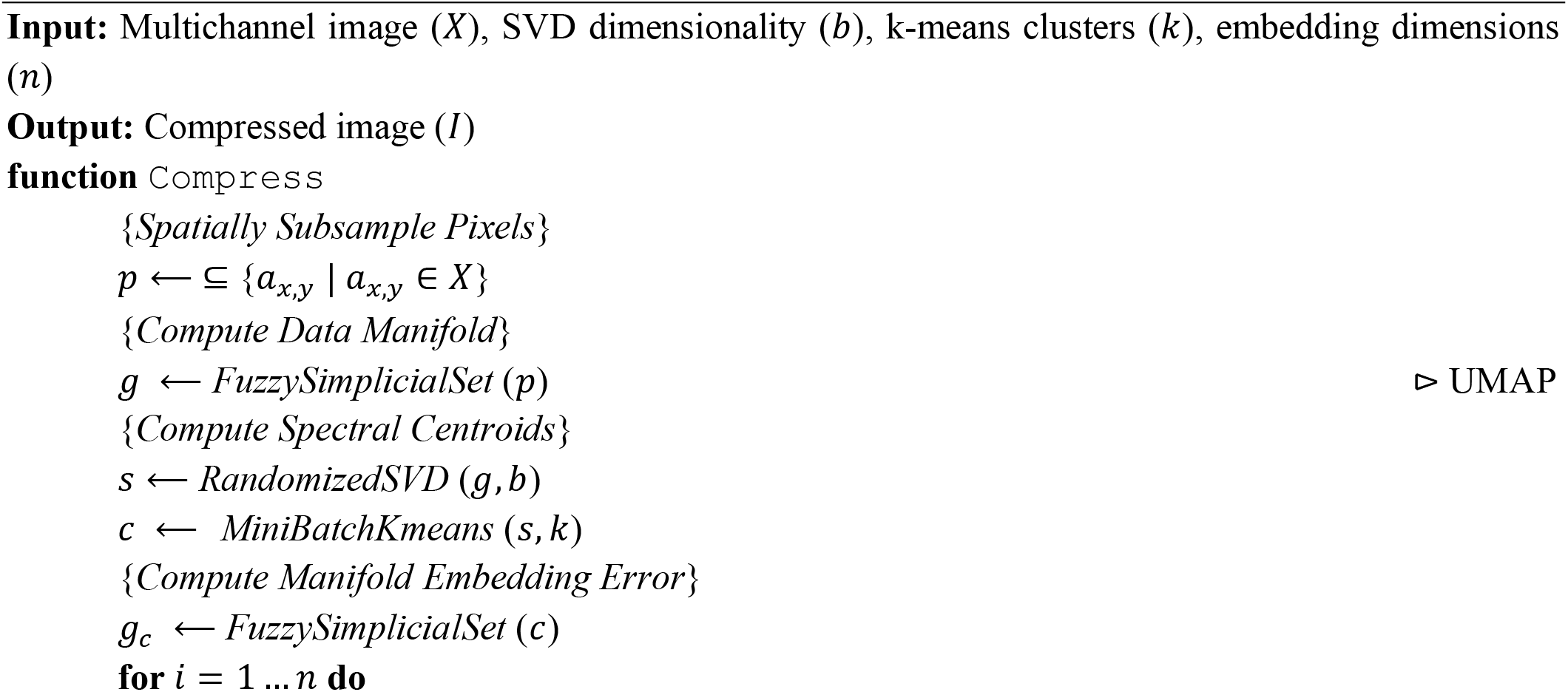
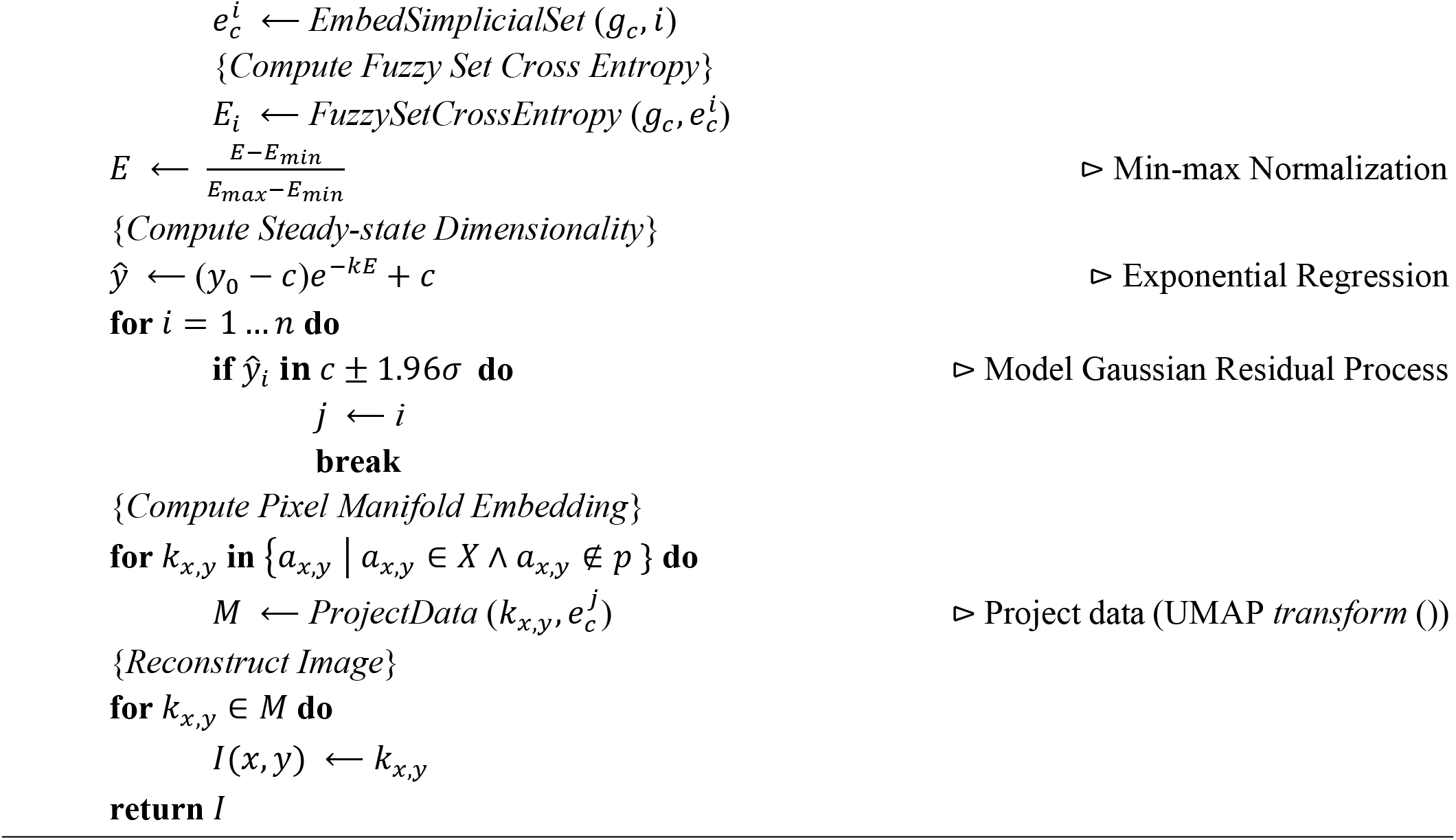

#### Image data subsampling

Subsampling is performed at pixel level and is optional for image compression. Implemented options include uniformly spaced grids within the (*x*, *y*) plane, random coordinate selection, and random selection initialized with uniformly spaced grids (“pseudo-random”). HDIprep also supports specification of masks for sampling regions, which may be useful for extremely large data sets.

By default, images with fewer than 50,000 pixels are not subsampled, images with 50,000-100,000 pixels are subsampled using 55% pseudo-random sampling initialized with 2×2 pixel uniformly spaced grids, images with 100,000-150,000 pixels are subsampled using 15% pseudo-random sampling initialized with 3×3 pixel grids, and images with more than 150,000 pixels are subsampled with 3×3 pixel grids. These default values are based on empirical studies (**Supplementary Figs. 4, 5, and 6**).

No subsampling was used for presented MSI data. Subsampling rates used for presented IMC data were determined on a case-by-case basis from empirical studies and match those used in the spectral landmark sampling experiments. Subsampling with 10×10 pixel uniformly spaced grids was used for CyCIF data compression.

#### Fuzzy simplicial set generation

To construct a pixel-level data manifold, we represent each pixel as a *d*-dimensional vector, where *d* is the number of channels in the given high-parameter image (i.e., discarding spatial information). We then implement the UMAP algorithm and extract the resulting fuzzy simplicial set representing the manifold structure of these *d*-dimensional points. For all presented results, we used the default UMAP parameters to generate this manifold: 15 nearest neighbors and the Euclidean metric.

#### Manifold landmark selection with spectral clustering

Spectral landmarks are identified using a variant of spectral clustering. We use randomized singular value decomposition (SVD) followed by mini-batch k-means to scale spectral clustering to large data sets, following the procedure introduced in the potential of heat diffusion for affinity-based transition embedding (PHATE) algorithm^18^. Given a symmetric adjacency matrix *A* representing pairwise similarities between nodes (here, pixels) originating from a *d*-dimensional space 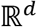, we first compute the eigenvectors corresponding to the *k* largest eigenvalues of *A*. We then perform mini-batch k-means on the nodes of *A* using these *k* eigenvectors as features. Spectral landmarks are defined as the d-dimensional centroids of the resulting clusters.

By default, the input data is reduced to 100 components using randomized SVD and then split into 3,000 clusters using mini-batch k-means. These default parameter values are based on empirical studies (**Supplementary Fig. 7**). Due to steady-state embeddings of MSI and IMC data only being available after experimental tests, no landmark selection was used for processing or determining the optimal embedding dimensionality of these data sets. Instead, full or subsampled datasets were used. All other steady-state embeddings for image data was compressed using the above default parameters.

#### Steady-state UMAP embedding dimensionalities

By default, HDIprep embeds spectral landmarks into Euclidean spaces with 1-10 dimensions to identify steady-state embedding dimensionalities. Exponential regressions on the spectral landmark fuzzy set cross entropy are performed using built-in functions from the Scipy Python library. These default parameters were used for all presented data.

#### Histology image preprocessing

HDIprep processing options for hematoxylin and eosin (H&E) stained tissues and other low-channel histological stains include image filters (e.g., median), thresholding (e.g., manually set or automated), and successive morphological operations (e.g., thresholding, opening and closing). Presented H&E and toluidine-blue stained images were processed using median filters to remove salt-and-pepper noise, followed by Otsu thresholding to create a binary mask representing the foreground. Sequential morphological operations were then applied to the mask, including morphological opening to remove small connected foreground components, morphological closing to fill small holes in foreground, and filling to close large holes in foreground.

#### Image compression with UMAP parametrized by a neuronal network

We implemented parametric UMAP^36^ using the default parameters and neural architecture with a TensorFlow backend. The default architecture was comprised of a 3-layer 100-neuron fully connected neuronal network. Training was performed using gradient descent with a batch size of 1,000 edges and the Adam optimizer with a learning rate of 0.001.

### High-dimensional image registration (HDIreg)

HDIreg is a containerized workflow implementing the open-source Elastix software^30^ in conjunction with custom-written Python modules to automate the image resizing, padding, and trimming often applied before registration. HDIreg incorporates several different registration parameters, cost functions, and deformation models, and additionally allows manual definition of point correspondences for difficult problems, as well as composition of transformations for fine-tuning (see **Supplemental Note 2**, **Notes on the HDIreg workflow’s expected performance**).

High-parameter images are registered using a manifold alignment scheme parametrized by spatial transformations, which aims to maximize image similarity. Formally, we view registration as the following optimization problem^40^:

Given a fixed *d*-dimensional image 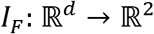 with domain Ω_*F*_ and a moving *q*-dimensional image 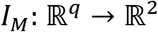 with domain Ω_*m*_, we aim to optimize

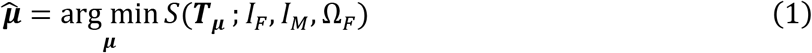

where ***T_μ_***: Ω_*F*_ → Ω_*M*_ is a smooth transformation defined by the vector of parameters 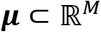, and *S* is a similarity measure maximized when *I_M_* ○ ***T_μ_*** and *I_F_* are aligned.

#### Differential geometry and manifold learning

MIAAIM’s manifold alignment scheme uses the entropic graph-based Rényi *α*-mutual information (*α*-MI)^40^ as the similarity measure S in **Equation 1**, which extends to manifold representations of images (i.e., compressed images) embedded in Euclidean space with potentially differing dimensionalities. This measure is justified in the HDIreg manifold alignment scheme through the notion of *intrinsic manifold information* (i.e., entropy). In what follows, we introduce basic differential geometric concepts that enable us to expand existing foundations of intrinsic manifold entropy estimation to the UMAP algorithm. We assume familiarity with the definition of a topological space (see Armstrong^46^ for an introduction).

##### Definition 2

Let *X* and *Y* be topological spaces. A function *f*:*X* → *Y* is *continuous* if for each point *x* ∈ *X* and each open neighborhood *N* of *f*(*x*) ⊆ *Y* the set *f*^-1^(*N*) is an open neighborhood of *x* ∈ *X*. A function *f*:*X* → *Y* is a *homeomorphism* if it is one-to-one, onto, continuous, and has a continuous inverse. When a homeomorphism exists between spaces *X* and *Y*, they are called *homeomorphic* spaces.

##### Definition 3.

A *manifold 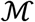 of dimension n* (i.e., an π-manifold) is a second-countable Hausdorff space, each point of which has an open neighborhood homeomorphic to *n*-dimensional Euclidean space, 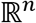. For any open set 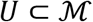 we can define a chart (*φ*, *U*) where 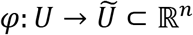 is a homeomorphism. We can say that (*φ*, *U*) acts as a local coordinate system for 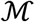, and we can define a transition between two charts (*φ*, *U*) and (*ω*, *V*) as *φ* ○ *ω*^-1^: *ω*(*U* ⋂ *V*) → *φ*(*U* ⋂ *V*) when *U* ⋂ *V* is non-empty.

##### Definition 4.

A *smooth manifold* is a manifold where there exists a smooth transition map between each chart of 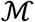. A *Riemannian metric g* is a mapping that associates to each point 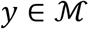 an inner product *g_y_*(·, ·) between vectors tangent to 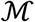 at *y*. We denote tangent spaces of *y* as 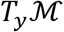. A *Riemannian manifold*, written 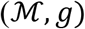, is a smooth manifold 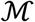 together with a Riemannian metric *g*. Given a Riemannian manifold, the *Riemannian volume element* provides a means to integrate a function with respect to volume in local coordinates. Given 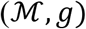, we can express the volume element *ω* in terms of the metric *g* and the local coordinates of a point *x* = *x*_1_,…, *x_n_* as *ω* = *g*(*x*)*dx*_1_ ∧… ∧ *dx_n_* where *g*(*x*) > 0 and ∧ indicates the wedge product. The volume of 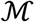 under this volume form is given by 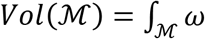.

##### Definition 5.

An *immersion* of a smooth *n*-manifold 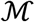 into 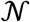 is a differentiable mapping 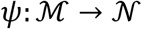 such that 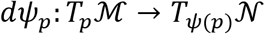 is injective for all points 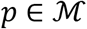. *ψ* is therefore an immersion if its derivative is everywhere injective.

##### Definition 6.

An *embedding* between smooth manifolds 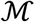 and 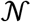 is a smooth function 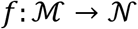 such that *f* is an immersion and an embedding of topological spaces (i.e., is an injective homeomorphism).

Let 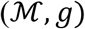 be a compact, *n*-dimensional Riemannian manifold immersed in an ambient 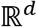, where *n* ≪ *d*, and let 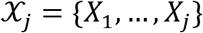 be a set of independent and identically distributed random vectors with values drawn from a distribution supported by 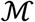. Define 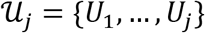 to be open neighborhoods of the elements of 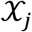. An aim of manifold learning is to approximate an embedding *f* such that a measure of distortion *D* is minimized between 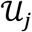 and 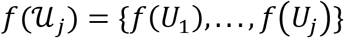. The manifold learning problem can therefore be written as

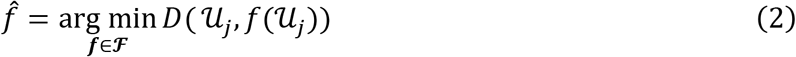

where 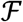 represents the family of possible measurable functions taking *x* ↦ *f*(*x*). In machine learning settings, open neighborhoods 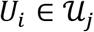 about vectors 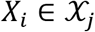 are often defined to be KNN neighborhoods with geodesic distances (or probabilistic encodings thereof^47^) within each open neighborhood approximated with a positive definite kernel^48^, which allows the computation of inner products in a Riemannian framework (as compared with a pseudo-Riemannian framework which need not be positive definite). Measures of distortion vary by algorithm (see **Supplementary Note 3, HDIprep dimension reduction validation** for examples). Of interest in our exposition are the measures induced by the embedded geodesics via the volume elements of these coordinate patches. These provide the components needed to quantify the *intrinsic Rényi α-entropy* of embedded data manifolds.

#### Entropic graph estimators

Given Lebesgue density *f* and identically distributed random vectors ***X***_1_,…, ***X**_n_* with values in a compact subset of 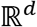, the *extrinsic* Rényi *α*-entropy of *f* is given by:

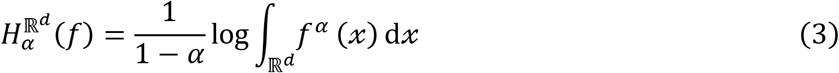

where *α* ∈ (0,1).

##### Definition 7 (adapted from Costa and Hero^38^)

Let 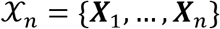 be identically distributed random vectors with values in a compact subset of 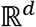, the nearest neighbor of 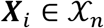 under the Euclidean metric is given by:

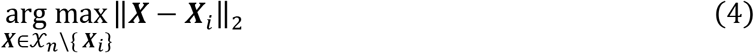

A *k*-nearest neighbor (KNN) graph puts and edge between each 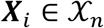 and its *k*-nearest neighbors. Let 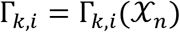 be the set of fc-nearest neighbors of 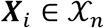. Then the total edge length of the KNN graph for 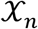 is given by:

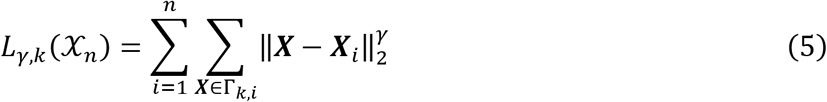

where *γ* > 0 is a power-weighting constant.

In practice, the *extrinsic* Rényi *α*-entropy of *f* can be suitably approximated using a class of graphs known as continuous quasi-additive graphs, including k-nearest neighbor (KNN) Euclidean graphs^49^, as their edge lengths asymptotically converge to the Rényi *α*-entropy of feature distributions as the number of feature vectors increases^50^. This property leads to the convergence of KNN Euclidean edge lengths to the *extrinsic* Rényi *α*-entropy of a set of random vectors with values in a compact subset of 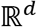 with *d* ≥ 2 ^38^. This is a direct corollary of the **Beardwood-Halton-Hammersley Theorem** outlined below.

### Beardwood-Halton-Hammersley (BHH) Theorem^38, 49^

*Let 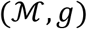 be a compact Riemannian m-manifold immersed in an ambient* 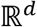. *Suppose 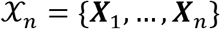 are identically distributed random vectors with values in a compact subset of* 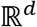 *and Lebesgue density f. Assume d* ≥ 2, 1 ≤ *γ* < *d and define 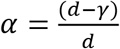. Then with probability 1*,

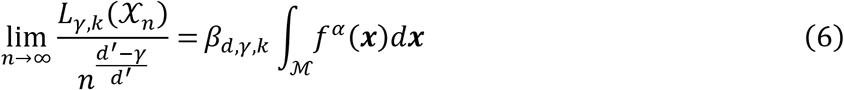

The value that determines the right side of the limit in **Equation 6** is the extrinsic Rényi *α*-entropy given by Equation 4. When identically distributed random vectors are restricted to a compact smooth *m*-manifold, 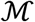, in an ambient 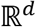, the **BHH Theorem** generalizes to enable an estimation of *intrinsic* Rényi *α*-entropy 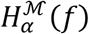 of the multivariate density *f* on 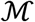 defined by

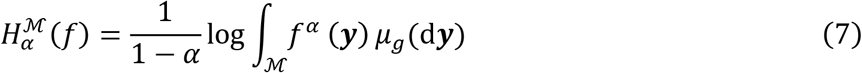

by incorporating the measure *μ_g_* naturally induced by the Riemannian metric via the Riemannian volume element^38^. This is formalized by the following given by Costa and Hero**^38^**:

#### Theorem 1 (Costa and Hero^38^)

*Let 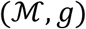 be a compact Riemannian m-manifold immersed in an ambient* 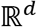. *Suppose 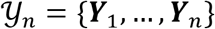 are identically distributed random vectors of 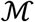 with bounded density f relative to the differential volume element μ_g_ induced by the metric μ_g_. Assume d* ≥ 2, 1 ≤ *γ* < *d and define 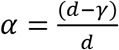. Then with probability 1*,

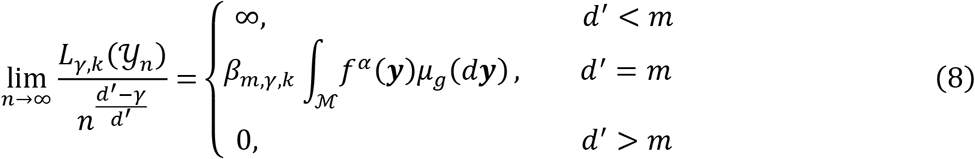

*where β_m,γ,k_ is a constant that is independent of f and 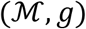. Similarly, the expectation 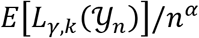 converges to the same limit*.

The quantity that determines the limit when *d*′ = *m* is the intrinsic Renyi alpha entropy of *f* given by **Equation 7. Theorem 1** has been used in conjunction with manifold learning algorithms Isomap and a variant C-Isomap to estimate the intrinsic dimensionality of embedded manifolds^39^. In contrast to these results that use all pairwise geodesic approximations for each point in the data set to estimate the *a*-entropy, we aim to provide a similar formulation utilizing local information contained in data manifolds, following the results of our dimension reduction benchmark which shows that local information preserving algorithms are well-suited for the task of high-dimensional image data compression (**Supplementary Figs 1, 2, 3**). Indeed, the *information density* of volumes of continuous regions of model families (i.e., collections of output embedding spaces or input points) have been recognized in defining an information geometry of statistical manifold learning^47^, motivating the title of our software.

#### Entropic graph estimators on local information of embedded manifolds

In what follows, we utilize two concepts to show that the intrinsic information of multivariate probability distributions supported by embedded manifolds in Euclidean space with the UMAP algorithm can be approximated using the **BHH Theorem**: (i.) the compactness of constructed manifolds and (ii.) the conservation of Riemannian volume elements. We address (i.) with a simple proof, and we provide a motivational example of conservation of volume elements using UMAP to address (ii.).

##### Definition 8.

A topological space *X* is *compact* if every open cover 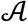 of *X* contains a finite sub collection that also covers *X*. By finite open cover, we mean that the elements of 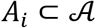 with *i* ∈ {1,…, *n*} are open sets in *X*, and that the union of the elements of 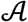 equals 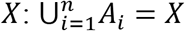.

##### Proposition 1.

*Let n > d and suppose that 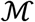 is a compact manifold of dimension r with r ≤ d that is immersed in ambient* 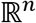. *Then the image 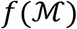 of 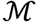 under a projection* 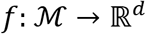 *is compact*.

*Proof*. Let 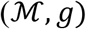 be a compact Riemannian manifold (e.g., a manifold constructed with UMAP) with metric *g* in an ambient 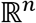 and *f* a projection from 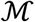 to 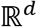. Since *f* is a projection, it is continuous, which takes compact sets to compact sets.

**Proposition 1** shows that a *d*-dimensional Euclidean projection of a compact Riemannian manifold take values in a compact subset of 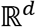, a necessary condition in the **BHH Theorem**. The UMAP algorithm considers fuzzy simplicial sets, i.e., manifolds, constructed from finite extended pseudo-metric spaces (see finite fuzzy realization functor, Definition 7, given by McInnes and Healy^35^). By finite, we mean that these extended pseudo-metric spaces are constructed from a finite collection of points. If one considers this finiteness condition, then the compactness of UMAP manifolds follows naturally from **Definition 8** – given an open cover on a manifold, one can find a finite subcover. Thus, UMAP projections are compact, following **Proposition 1**.

To extend the **BHH Theorem** to the calculation of intrinsic *α*-entropy of UMAP embeddings as in **Equation 7**, we must show that volume elements induced via embedding are well approximated. Note that these results apply to any dimension reduction algorithm that can provably preserve distances within open neighborhoods upon embedding a compact manifold in Euclidean space. In what follows, we do not provide a proof that UMAP preserves distances within open neighborhoods about points, although, this would be an ideal scenario. Rather, we assume that this ideal scenario exists, and we describe how to find the optimal dimensionality for projecting data to satisfy this assumption.

In contrast to global data preserving algorithms, such as Isomap that calculate all pairwise geodesic distances or approximations thereof with landmark based approaches, UMAP approximates geodesic distances in open neighborhoods local to each point (see **Lemma 2** below). Given Lebesgue density *μ* and identically distributed random vectors ***Y***_1_,…, ***Y***_n_ with values constrained to lie on a compact, uniformly distributed Riemannian manifold 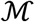, geodesics between samples ***Y**_i_* and ***Y**_j_* drawn from *μ* are encoded probabilistically with UMAP and represent a scaled exponential distribution^51^:

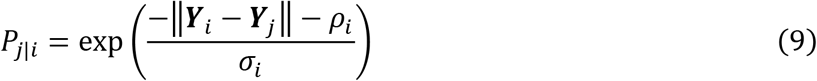

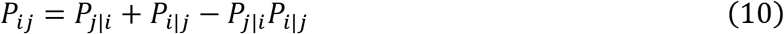

where *ρ_i_* is the distance from vector ***Y**_i_* to its nearest neighbor and *σ_i_* is an adaptively chosen normalization factor. Using the terminology in **Equation 2**, the objective of embedding in UMAP is given by minimizing the fuzzy simplicial set cross-entropy (**Definition 1**), which represents distortion *D*. Formally, given probability distribution *P_ij_* encoding geodesics between samples ***Y**_i_* and *Y_j_* and probability distribution *Q_ij_* encoding distances between samples *f*(***Y**_i_*) and *f*(***Y**_j_*), we can represent the cross-entropy loss employed by UMAP as (adapted from Narayan et. al^51^):

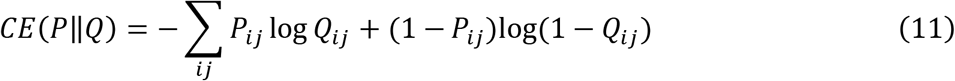

where *Q_ij_* is the probability distribution formed from low dimensional positions of embedded vectors *f*(***Y**_i_*) and *f*(***Y**_j_*) by 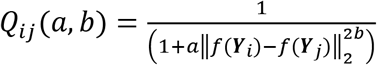 with *a*,*b* user-defined parameters to control embedding spread.

Minimizing **Equation 11** is not, in general, a convex optimization problem. Optimization over the family 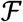 from **Equation 2** is restricted to a subset rather than the full family and thus represents, in the best case, a local optimum. We include a larger family of measurable functions in order to more accurately approach optimal embedding of geodesic distances within open neighborhoods of vectors through the identification steady-state embedding dimensionalities, as outlined in the HDIprep workflow in a “pseudo-global” optimization procedure.

Using the notation in **Equation 2**, the volume elements induced by the embedding 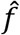 are similarly those that minimize distortion between the volume elements of 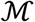 given for local coordinates *x*_1_,…,*x_n_* by *σ* = *g*(*x*)*dx* where *dx* = *dx*_1_ ∧… ∧ *dx_n_* and those of 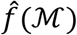 given for local coordinates 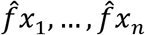 by 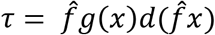, assuming that the dimensionality *n* of local coordinates is preserved. Under a globally optimal solution to **Equation 7 (** i.e., when *P_ij_* = *Q_ij_*), volume elements are preserved: 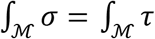. The existence of volume preserving diffeomorphisms for compact manifolds in consideration are proven by Moser^52^.

In order to identify manifold embeddings that minimize distortion of volume elements, we viewed increases in the dimensionality of real-valued data in a natural way by modeling increases in dimensionality as exponential increases in potential positions of points (i.e., increasing copies of the real line, 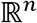). The relationship between radii of open neighborhoods, volume, and density of manifolds in a learning setting is formalized by Narayan et. al^51^ in an application where density is preserved with a given dimensionality for embeddings by altering open neighborhood radii; however, we can readily extend this in an adapted scenario of volume preservation under fixed radii to infer a dimensionality that satisfies the assumption of geodesic distance preservation. Consider the following example:

Let 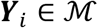 be a vector of manifold 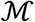 immersed in an ambient 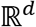, and assume that the *k*-nearest neighbors of ***Y**_i_* are uniformly distributed in a ball *B_r_d__* of radius *r_d_* with proportional volume 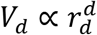. Assume that a map *f* takes the open neighborhood *B_r_d__* of ***Y**_i_* to an *m*-dimensional ball 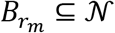 of manifold 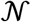 with radius *r_m_* while preserving its structure, including the uniform distribution and the induced Riemannian volume element *V_m_*. Following Narayan et al.^51^, we can use the proportion 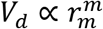 to infer a power law relationship 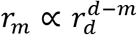 between the local radii *r_m_* in embedding spaces and the original radius of *B_r_d__*.

To extend this example, suppose that *r_m_* and *r_d_* are fixed (we can assume that radii are fixed in the original space, and that those in the embedding space are controlled by the *a*, *b* parameters influencing *Q_ij_* in **Equation 11**). Assume that the ambient metrics of the embedding space and the original space are the same, and that they give rise to geodesic distances within *B_r_m__* and *B_r_d__* using the native UMAP method (**Lemma 2** below). Since ambient metrics and radii are preserved and that 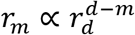, this implies that the geodesic distances *δ_m_* and *δ_d_* between points in *B_r_m__* and *B_r_d__* also exhibit a power law relationship, 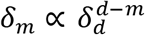. If we further assume that 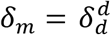 (i.e., that geodesic distances between points in open neighborhoods are preserved, the ideal scenario) we can solve for the relationship between geodesics and dimensionality *m*. Specifically, suppose that 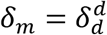. Substituting 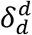 for *δ_m_*, we can see that

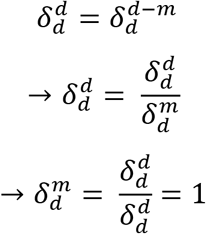

which implies that the geodesics within open neighborhoods of points of the original space have an exponential relationship with dimensionality *m*.

Using the power-law relationship between volumes of open neighborhoods in UMAP manifolds and their embedded counterparts, we can attempt to identify the dimensionality *m* that such that geodesics within open neighborhoods is preserved with exponential regression. Note that geodesic distance preservation is stronger than volume preservation; however, volume preservation is implied. As a result, steady-state manifold embeddings provide a Euclidean dimensionality to approximate manifold geodesics of vectors sampled from 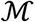, and thus the volume elements of 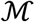. KNN graph functionals calculated in the steady-state embedding space provide the necessary machinery to calculate the intrinsic *α*-entropy of embedded data manifolds in MIAAIM by applying the **BHH Theorem** using the induced measure across all coordinate patches as in **Theorem 1**.

Note, though, that the distances outside of open neighborhoods of points are not guaranteed to be accurately modelled in the embedding space with UMAP if one makes the assumptions introduced in our example. Therefore, applying **Theorem 1** in conjunction with UMAP by replacing KNN graph lengths with those obtained with length functionals of geodesic minimal spanning trees (GMST), another type of entropic graph, should not be expected to reproduce the intrinsic entropy originally reported by Costa and Hero^39^. Our primary contribution here is combining the KNN intrinsic entropy estimator with a local information preserving dimension reduction algorithm. We wish to extend these results to the setting where two such manifolds are compared, as this serves the basis for our image registration application. An entropic graph-based estimator of *α*-MI in an image registration setting is described by the following^40^:

Let *z*(*x_i_*) = [*Z_i_*(*x_i_*),…, *z_d_*(*x_i_*)] be a d-dimensional vector encoding the features of point *x_i_*. Let *Z_f_*(*x*) = {*z^f^*(*x*_1_),…, *z^f^*(*x_N_*)} be the feature set of a fixed image, *Z_m_*[***T_μ_***(*x*)) = {*z^m^*(***T_μ_***(*x*_1_)),…, *z^m^*(***T_μ_***(*x_N_*))} be the feature set of a transformed moving image at points in *T_μ_*(*x*), and 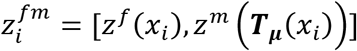 be the concatenation of the feature vectors of the fixed and transformed moving image at *x_i_*. Then,

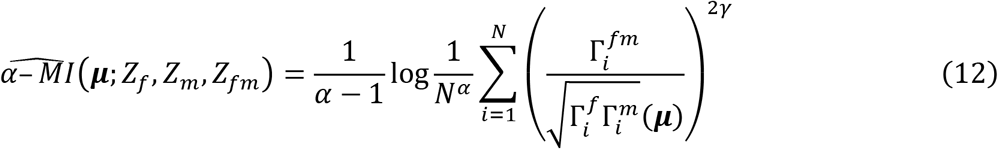

is a graph-based estimator for *α*-MI, where *γ* = *d*(1 – *α*), 0 < *α* < 1, and three graphs

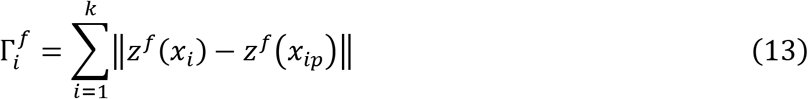

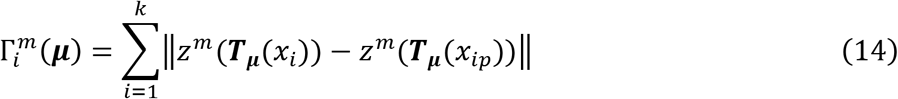

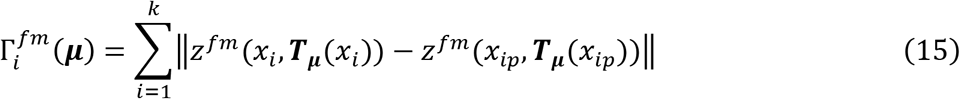

represent the Euclidean graph functionals (lengths) of a feature vector *z* to its *p^th^* nearest neighbor over *k* considered nearest neighbors.

The Rényi *α*-MI provides a quantitative measure of association between the intrinsic structure of multiple manifold embeddings constructed with the UMAP algorithm. The Rényi *α*-MI measure extends to feature spaces of arbitrary dimensionality, which MIAAIM utilizes in combination with its image compression method to quantify similarity between steady-state embeddings of image pixels in potentially differing dimensionalities.

#### Multimodal image registration studies

Since data acquisition removed tissue context at regions of interest on the IMC full-tissue reference image, we first aligned full tissue sections then used the coordinates of IMC regions to extract data from all modalities for fine-tuning. Custom Python scripts were used to propagate alignment across imaging modalities. Manual landmark correspondences were used where unsupervised alignment proved suboptimal. We accounted for alignment errors around IMC regions following full-tissue registration by padding regions with extra pixels prior to cropping. All registrations involving MSI or IMC data were conducted using KNN *α*-MI, as in **Equation 12**, with *α* = 0.99 and 15 nearest neighbors. All registrations aligning low-channel slides (IMC reference toluidine blue image and H&E) were conducted using histogram-based MI after grayscale conversion for rapid processing.

For full-tissue images, a two-step registration process was implemented by first aligning images using an affine model for the vector of parameters ***μ* (Equation 1**) and subsequently aligning images with a nonlinear model parametrized by B-splines. Hierarchical Gaussian smoothing pyramids were used to account for resolution differences between image modalities, and stochastic gradient descent with random coordinate sampling was used for optimization. We additionally optimized final control point grid spacings for B-spline models and the number of hierarchical levels to include in pyramidal smoothing for each MSI data set to H&E alignment individually (**Supplementary Figs. 1, 2, 3**). A final control point spacing of 300 pixels for nonlinear B-spline registrations of MSI data to corresponding H&E data balanced correct alignment with unrealistic warping, which we identified visually and by inspecting the determinants of the spatial Jacobian matrices for values that deviated substantially from 1. H&E and IMC reference tissue registrations utilized a final grid spacing of 5 pixels. Similar optimizations for numbers of pyramidal levels were carried out for these data. All data that underwent image registration were exported and stored as 32-bit NIfTI-1 images. IMC data was not transformed and was kept in 16-bit OME-TIF(F) format.

### Cobordism Approximation and Projection (PatchMAP)

PatchMAP is an algorithm that constructs a smooth manifold by representing lower dimensional manifolds as its boundary, and it projects the higher-order manifold in a lower dimensional space for visualization. Higher-order manifolds produced by PatchMAP can be understood as cobordisms, which are described by the following set of definitions:

#### Definition 9.

Given a family of sets {*S_i_*: *i* ∈ *I*} and an index set *I*, their *disjoint union*, denoted *S* = ∐_*i*∈*I*_*S_i_* is a set together with injective functions *φ_i_*: *S_i_* → *S* for each *S_i_*. The disjoint union corresponds to the coproduct of sets.

#### Definition 10.

Two closed *n*-manifolds 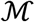 and 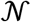 are *cobordant* if their disjoint union, denoted 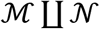, is the boundary of an *n*+1-manifold 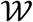. We call the manifold 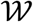 a *cobordism*. By boundary of a *d*-manifold, we mean the set of points on 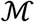 which are homeomorphic to the upper half plane, 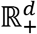. We denote the boundary of 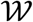 as 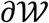.

PatchMAP addresses cobordism learning in a semi-supervised manner, where data is assumed to follow the structure of a nonlinear cobordism, and our task is to glue lower dimensional manifolds to the boundary of a higher-dimensional manifold to produce a cobordism. Here, we would like for coordinate transformations across the cobordism to have their own geometries independent of the metrics of boundary manifolds. In practice, this property allows us to explore data within boundary manifolds without being reliant on the specific geometry of the cobordism. Ultimately, cobordism geodesics are the base component of downstream applications such as the i-PatchMAP workflow. Furthermore, we would like for the cobordism to highlight boundary manifolds whose data points are highly similar. A natural way to satisfy both of these conditions is to use the fuzzy set-theoretic foundation of the UMAP algorithm.

The primary goal of PatchMAP then is to identify a smooth manifold whose boundary is the disjoint union of smooth manifolds of lower dimensionality, and which has a metric independent of each boundary manifolds’ metric that we choose to represent. We address this with a two-step algorithm that uncouples the computation of boundary manifolds from the cobordism. First, boundary manifolds are computed by applying the UMAP algorithm to each set of data with a user-provided metric. Practically, the result of this step are symmetric, weighted graphs that represent geodesics within each boundary manifold. Our task is to construct from a finite set of *n* boundary manifolds 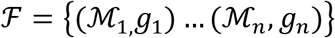 a manifold 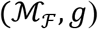, with metric *g* such that the disjoint union 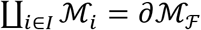. We wish to approximate the geodesics of 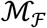, and for each element of 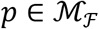, the tangent space 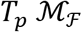 with inner product *g_p_*.

#### Lemma 2 (McInnes and Healy^35^)

*Let 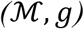 be a Riemannian manifold in an ambient* 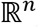. *and let 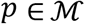 be a point. If g is locally constant about p in an open neighborhood 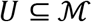 such that g is a constant diagonal matrix in ambient coordinates, then in a ball B ⊆ U centered at p with volume 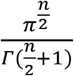 with respect to g, the geodesic distance from p to any point in q ∈ B is 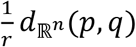, where r is the radius of the ball in the ambient space 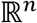 and 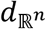 is the metric on the ambient space*.

If we assume that data points across boundary manifolds can be compared with a suitable metric, we can compute the geodesic distances between points on the projections of each boundary manifold under a user-provided ambient metric using **Lemma 2** (for a proof of **Lemma 2**, see Appendix A of McInnes and Healy^35^) to their disjoint union. Given two boundary manifolds 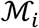 and 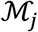, we compute the pairwise geodesic distances between points *p_i_*, 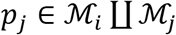 where 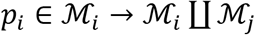 and 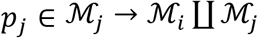 to obtain the components needed to construct the metric on 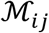 where the boundary 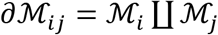. By extension, concatenation of all pairwise distance calculations between boundary manifolds 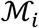 and 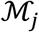 where *i*, *j* ∈ {1,2,…,*n*}; *i* ≠ *j* to their disjoint union 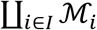 provides the components to construct the geodesics on the full cobordism 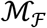 across all pairwise combinations of boundary manifolds, as desired. The result of using **Lemma 2** to approximate the projections of manifold geodesics, however, is that we have directed, incompatible views of the geodesics across the cobordism, to and from, boundary manifolds. That is, 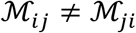. We can interpret these directed geodesics as being defined on *oriented* cobordisms. We aim to encode the directed geodesics in the oriented cobordisms and the boundary manifold geodesics with a single data representation.

Our goal then is to construct an *unoriented* cobordism, where the directed geodesics of oriented cobordisms are resolved into a single, symmetric matrix representation. To do this, we can translate each of the extended pseudo-metric spaces described above into a fuzzy simplicial set using the fuzzy singular set functor (see Definition 9 by McInnes and Healy^35^), which captures both a topological representation of the oriented cobordisms and the underlying metric information. The incompatibilities in oriented cobordism geodesics can be resolved with a norm of our choosing. A natural choice for the fuzzy set representation that we have chosen is the t-norm (also known as a *fuzzy intersection).* If we interpret the fuzzy simplicial set representations of oriented cobordisms in a probabilistic manner, then their intersection corresponds to the joint distribution of the oriented cobordism metric spaces, which highlights the directed cobordism geodesics that occur in both directions, to and from, with a high probability.

After symmetrizing the cobordisms between each pairwise boundary manifold, the final step is to integrate the boundary manifold geodesics with the symmetric cobordism geodesics obtained with the fuzzy set intersection. Additionally, we should ensure that cobordisms with shared domains (i.e., boundary manifolds) are identified and integrated with each other. We can do this by taking a *fuzzy set union* (probabilistic t-conorm) over the family of extended pseudo-metric spaces, as in the original UMAP implementation. The result is a cobordism that contains its own geometric structure that is captured in cobordism geodesics, in addition to individual boundary manifolds that contain their own geometries.

Optimizing a low dimensional representation of the cobordism can be achieved with a number of methods – we choose to optimize the embedding using the fuzzy set cross entropy (**Definition 1**), as in the original UMAP implementation for consistency. Note that since our algorithm produces a symmetric matrix, PatchMAP could be repeatedly applied to construct “nested” cobordisms of hierarchical dimensionality.

#### PatchMAP implementation

To construct cobordisms, PatchMAP first computes boundary manifolds by constructing fuzzy simplicial sets from each provided set of data, i.e., tissue state, by applying the UMAP algorithm (*FuzzySimplicialSet*, **Algorithm 2**). Then, pairwise, directed nearest neighbor (NN) queries between boundary manifolds are computed in the ambient space of the cobordism (*DirectedGeodesics*, **Algorithm 2**). Directed NN queries between boundary manifolds are weighted according to the native implementation in UMAP, the method of which we refer the reader to **Equations 5 and 6**. Resulting pairwise directed NN graphs between UMAP boundary manifolds are weighted, and they reflect Riemannian metrics that are not compatible. That is, they cannot be simply added or multiplied to integrate their weights. We therefore stitch the cobordism metric and make directed NN queries compatible by applying a fuzzy simplicial intersection, which results in a weighted, symmetric graph (*FuzzyIntersection*, **Algorithm 2**). The final cobordism produced by PatchMAP is obtained by taking a fuzzy union over the family of all fuzzy simplicial sets that represent cobordisms between boundary manifolds (*FuzzyUnion*, **Algorithm 2**). To represent connections between boundary manifolds in PatchMAP cobordism projections, we implemented the hammer edge bundling algorithm in the Datashader Python library. Pseudo-code outlining the PatchMAP algorithm is shown in the following:

#### Algorithm 2

PatchMAP

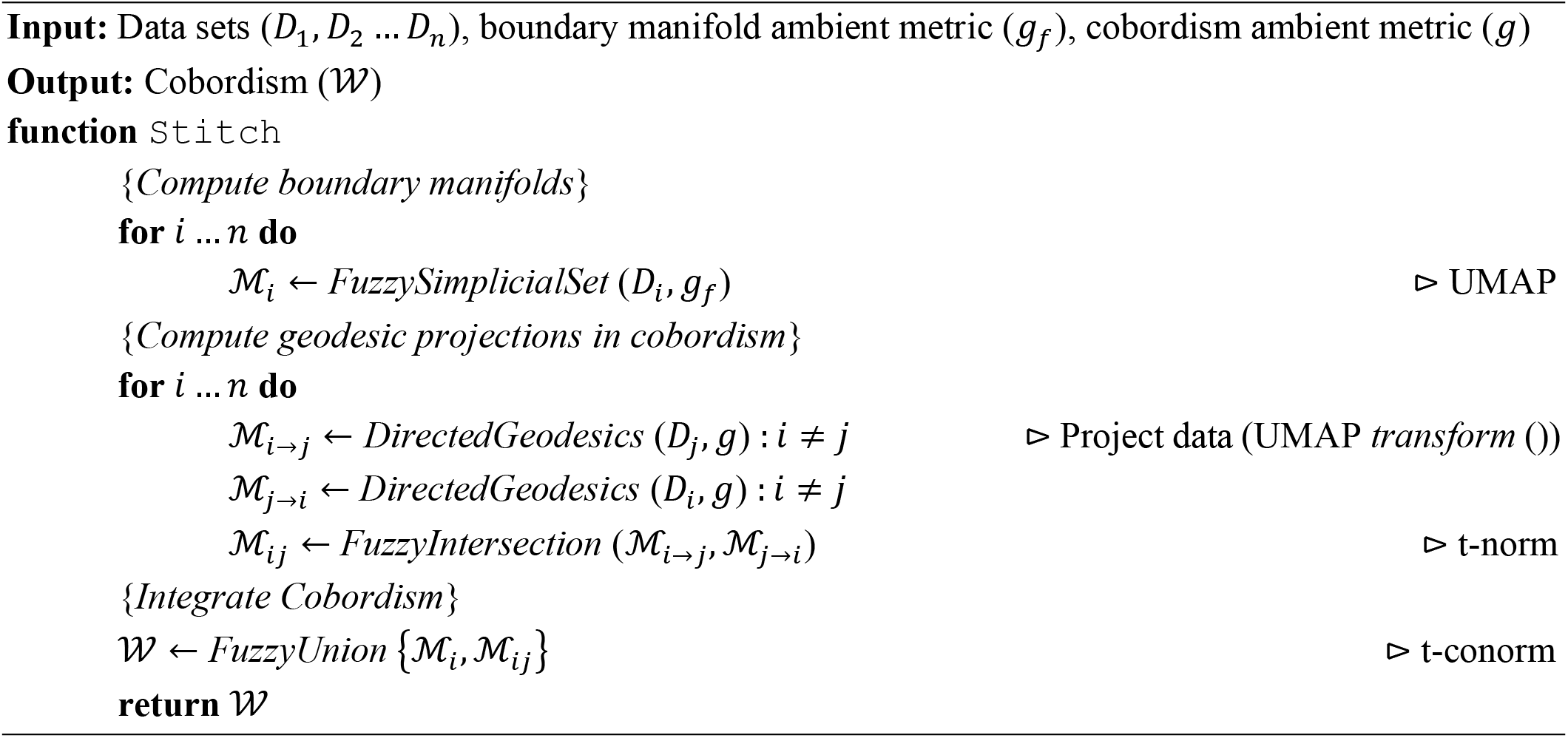

### Domain/Information Transfer (i-PatchMAP)

Let 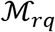 be a cobordism with boundary manifolds 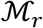 and 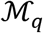 obtained with PatchMAP that come a reference and query data set, respectively. Specifically, 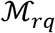 is a matrix where rows represent points in the reference boundary manifold, columns represent the nearest neighbors of reference manifold points in the query boundary manifold under a user defined metric, and the *i*, *j*^th^ entry represents the geodesics between points *p_i_*, 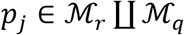 such that 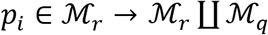 and 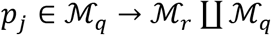. We compute a new feature matrix of predictions *P_q_* for the query data set by multiplying the feature matrix to be transferred, *F*, with the transpose of the weight matrix *W_rq_* obtained through *ℓ*_1_ normalization of 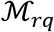:

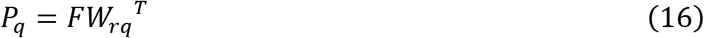

In this context, the matrix *W_rq_* can be interpreted as a single-step transition matrix of a Markov chain between states *p_i_* and *p_j_* derived from geodesic distances on the cobordism 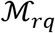.

### Biological Methods

All patient tissue samples were obtained with approval from the Institutional Review Boards (IRB) of Massachusetts General Hospital (protocol #2005P000774) and Beth Israel Deaconess Medical Center (protocol #2018P000581).

#### Generation of imaging mass cytometry data

Frozen tissues were sectioned serially at a thickness of 10 *μ*m using a Microm HM550 cryostat (Thermo Scientific) and thaw-mounted onto SuperFrost^™^ Plus Gold charged microscopy slides (Fisher Scientific). After temperature equilibration to room temperature, tissue sections were fixed in 4% paraformaldehyde (Ted Pella) for 10 min, then rinsed 3 times with cytometry-grade phosphate-buffered saline (PBS) (Fluidigm). Unspecific binding sites were blocked using 5% bovine serum albumin (BSA) (Sigma Aldrich) in PBS including 0.3% Triton X-100 (Thermo Scientific) for 1 hour at room temperature. Metal conjugated primary antibodies (Fluidigm) at appropriately titrated concentrations were mixed in 0.5% BSA in DPBS and applied overnight at 4 °C in a humid chamber. Sections were then washed twice with PBS containing 0.1% Triton X-100 and counterstained with iridium (Ir) intercalator (Fluidigm) at 1:400 in PBS for 30 min at room temperature. Slides were rinsed in cytometry-grade water (Fluidigm) for 5 min and allowed to air dry. Data acquisition was performed using a Hyperion Imaging System (Fluidigm) and CyTOF Software (Fluidigm), in 33 channels, at a frequency of 200 pixels/second and with a spatial resolution of 1 *μ*m. Images were visualized with MCD Viewer software (Fluidigm) before exporting the data as text files for further analysis. After imaging, slides were rapidly stained with 0.1% toluidine blue solution (Electron Microscopy Sciences) to reveal gross morphology. Slides were digitized at a resolution of approximately 2.75 *μ*m/pixel using a digital camera.

#### Generation of mass spectrometry imaging data

Paired 10*μ*m thick sections from the same tissue blocks as used for imaging mass cytometry were thaw mounted onto Indium-Tin-Oxide (ITO) coated glass slides (Bruker Daltonics). Tissue sections were coated with 2.5-dihydroxybenzoic acid (40 mg/mL in 50:50 acetonitrile:water including 0.1% TFA) with an automated matrix applicator (TM-sprayer, HTX imaging). Mass spectrometry imaging of sections was performed using a rapifleX MALDI Tissuetyper (Bruker Daltonics, Billerica, MA). Data acquisition was performed using FlexControl software (Bruker Daltonics, Version 4.0) with the following parameters: positive ion polarity, mass scan range (*m/z)* of 300–1000, 1.25 GHz digitizer, 50 *μ*m spatial resolution, 100 shots per pixel, and 10 kHz laser frequency. Regions of interest for data acquisition were defined using FlexImaging software (Bruker Daltonics, version 5.0), and individual images were visualized using both FlexImaging and SCiLS Lab (Bruker Daltonics). After data acquisition, sections were washed with PBS and subjected to standard hematoxylin and eosin histological staining followed by dehydration in graded alcohols and xylene. The stained tissue was digitized at a resolution of 0.5 *μ*m/pixel using an Aperio ScanScope XT brightfield scanner (Leica Biosystems).

### Mass spectrometry imaging data preprocessing

Data were processed in SCiLS LAB 2018 using total ion count normalization on the mean spectra and peak centroiding with an interval width of ±25mDa. For all analyses, a peak range of *m/z* 400-1,000 was used after peak centroiding, which resulted in 9,753 *m/z* peaks. No peak-picking was performed for presented data unless explicitly stated. Data were exported from SCiLS Lab as imzML files for further analysis and processing.

### Single-cell segmentation

To quantify parameters of single-cells in IMC and registered MSI data within the DFU dataset, we performed cell segmentation on IMC ROIs using the pixel classification module in Ilastik (version 1.3.2)^41^, which utilizes a random forest classifier for semantic segmentation. For each ROI, two 250 *μ*m by 250 *μ*m areas were cropped from IMC data and exported in the HDF5 format to use for supervised training. To ensure each cropped area was a representative training sample, a global threshold for each was created using Otsu thresholding on the Iridium (nuclear) stain with Scikit-image Python library. Cropped regions were required to contain greater than 30% of pixels over their respective threshold.

Training regions were annotated for “background”, “membrane”, “nuclei”, and “noise”. Random forest classification incorporated Gaussian smoothing features, edges features, including Laplacian of Gaussian features, Gaussian gradient magnitude features, and difference of Gaussian features, and texture features, including structure tensor eigenvalues and Hessian of Gaussian eigenvalues. The trained classifier was used to predict each pixels’ probability of assignment to the four classes in the full images, and predictions were exported as 16-bit TIFF stacks. To remove artifacts in cell staining, noise prediction channels were Gaussian blurred with a sigma of 2 and Otsu thresholding with a correction factor of 1.3 was applied, which created a binary mask separating foreground (high pixel probability to be noise) from background (low pixel probability to be noise). The noise mask was used to assign zero values in the other three probability channels from Ilastik (nuclei, membrane, background) to all pixels that were considered foreground in the noise channel. Noise-removed, three-channel probability images of nuclei, membrane, and background were used for single-cell segmentation in CellProfiler (version 3.1.8)^53^.

### Single-cell parameter quantification

Single-cell parameter quantification for IMC and MSI data were performed using an in-house modification of the quantification (MCQuant) module in the multiple-choice microscopy software (MCMICRO)^24^ to accept NIfFTI-1 files after cell segmentation. IMC single-cell measures were transformed using 99^th^ percentile quantile normalization prior to downstream analysis.

### Imaging mass cytometry cluster analysis

Cluster analysis was performed in Python using the Leiden community detection algorithm with the leidenalg Python package. UMAP’s simplicial set (weighted, undirected graph) created with 15 nearest neighbors and Euclidean metric was used as input to community detection.

### Microenvironmental correlation network analysis

To calculate associations across MSI and IMC modalities, we used Spearman’s correlation coefficient in the Python Scipy library. *M/z* peaks from MSI data with no correlations to IMC data with Bonferroni corrected P-values above 0.001 were removed from the analysis. Correlation modules were formed with hierarchical Louvain community detection using the Scikit-network package. The resolution parameter used for community detection was chosen based on the elbow point of a graph plotting resolution vs. modularity of community detection results. UMAP’s simplicial set, created with 5 nearest neighbors and the Euclidean metric, was used as input for community detection after inverse cosine transformation of Spearman’s correlation coefficients to form metric distances^54^. Visualization of MSI correlation module trends to IMC parameters were computed using exponential-weighted moving averages in the Pandas library in Python after standard scaling IMC and MSI single-cell data. MSI moving averages were additionally min-max scaled to a range of 0-1 for plotting purposes. Differential correlations of variables *u* from MSI data and *v* from IMC data between conditions *a* and *b* were quantified and ranked using the formula given by Hsu et al.^55^:

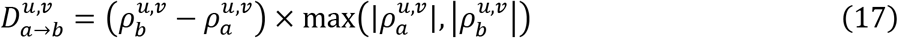

where change in correlation coefficients for each pair *u*, *v* between conditions are weighted according to the maximal absolute correlation coefficient among both conditions. Significance of differential correlations 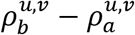 were calculated using one-sided, Bonferroni corrected z-statistics after Fisher transformation.

### Dimensionality reduction algorithm benchmarking

Methods used for benchmarking dimensionality reduction algorithms are outlined in **Supplementary Note 3, HDIprep dimension reduction validation**.

### Spatial subsampling benchmarking

Default subsampling parameters in MIAAIM are based on experiments across IMC data from DFU, tonsil, and prostate cancer tissues recording Procrustes transformation sum of squares errors between subsampled UMAP embeddings with subsequent projection of out-of-sample pixels and full UMAP embeddings using all pixels. Spatial subsampling benchmarking was performed across a range of subsampling percentages.

### Spectral landmark benchmarking

Subsampling percentages and dimensionalities to validate landmark-based steady-state UMAP dimensionalities were determined on a case-by-case basis from empirical studies and match those used for presented, registered data. Parameters were chosen due to the computational burden of computing values for cross-entropy on large data. To compare landmark steady-state dimensionality selections to subsampled data, we compared the shape of exponential regression fits from both sets of data using sum of squared errors. Sum of squared errors were calculated across a range of resulting landmarks.

### Cobordism boundary manifold stitching simulation

Simulations were performed using the MNIST digits dataset in the Python Scikit-learn library using the default parameters for BKNN, Seurat v3, Scanorama, and PatchMAP across a range of nearest neighbor values. Data points were split into according to their digit label and stitched together using each method. Integrated data from each tested method excluding PatchMAP was then visualized with UMAP. Quality of boundary manifold stitching for each algorithm was quantified using the silhouette coefficient in the UMAP embedding space, implemented in Python with the Scikit-learn library. The silhouette coefficient is a measure of dispersion for a partition of a dataset. A high value indicates that data from the same label/type are tightly grouped together, whereas a lower value indicates that data from different types are grouped together. The silhouette coefficient (SC) is the average silhouette score s computed across each data point in the dataset, given by the following:

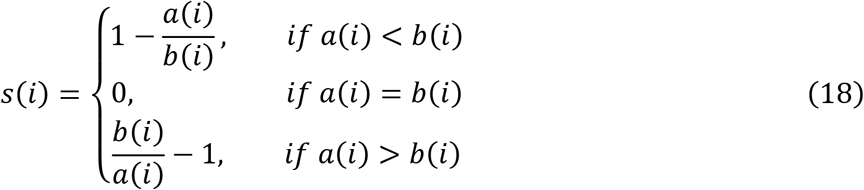

where *a*(*i*) is the average distance of data point *i* to all points with its label and *b*(*i*) is the average distance of point *i* to all other data that do not have the same label.

### CBMC CITE-seq data transfer

CBMC CITE-seq data were preprocessed according to the vignette provided by the Satija lab at https://satijalab.org/seurat/articles/multimodal_vignette.html. RNA profiles were log transformed and ADT abundances were normalized using centered log ratio transformation. RNA variable features were then identified, and the dimensionality of the RNA profiles of cells were reduced using principal components analysis. The first 30 principal components of single-cell RNA profiles were used to predict single-cell ADT abundances. The CBMC dataset was split randomly into 15 evaluation instances with 75% training and 25% testing data. Training data was used to predict test data measures. Prediction quality was quantified using Pearson’s correlation coefficient between true and predicted ADT abundances. Correlations were calculated using the Python library, Scipy. Seurat was implemented using the TransferData function after finding transfer anchors (FindTransferAnchors function) using the default parameters. PatchMAP and UMAP+ were applied with 80 nearest neighbors and the Euclidean metric in the PCA space.

### Spatially resolved image data transfer

To benchmark MSI to IMC information transfer, we performed leave-one-out cross validation with segmented single-cells from 23 image tiles (number of cells ranging from ~100 to ~500 each) from the DFU dataset. IMC ROIs were split into four evenly sized quadrants to create 24 tiles. One tile was removed due to lack of cell content. Data was transformed prior to information transfer using principal components analysis with 15 components using the Scikit-learn library. Seurat was implemented using the TransferData function after finding transfer anchors (FindTransferAnchors function) using the default parameters and 15 principal components. PatchMAP and UMAP+ were implemented using 80 nearest neighbors and the Euclidean metric in the PCA space. Information transfer quality was computed for each predicted IMC parameter by calculating Pearson’s correlation in Python with the Scipy library between the Moran’s autocorrelation index of ground truth data and predicted data. Moran’s autocorrelation index (*I*) is given by the following^13^:

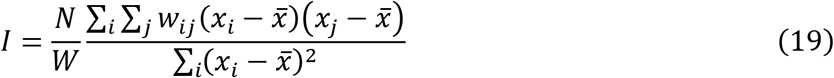

where *N* is the number of spatial dimensions in the data (2 for our purposes), *x* is the abundance of a protein of interest, 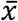 is the mean abundance of protein *x*, *w_ij_* is a spatial weight matrix, and *W* is the sum of all *W_ij_*.

We acknowledge the Satija lab for utilizing spatial autocorrelation to quantify information transfer quality in spatial data^13^. We calculated Moran’s I using the R package ape. Spatial weight matrices were calculated using the R package stats. To account for resolution differences between IMC and MSI imaging modalities, spatial weights for predicted IMC parameters were calculated using 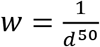 and weights for true IMC parameters were calculated using 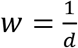 where *d* is the Euclidean distance between segmented cell centroids in the tiled image. Since there was no spatial deconvolution for spatial parameter predictions, IMC predictions were at a lower resolution than the true IMC data. Thus, a value of 50 was chosen to reduce the spatial weighting of predicted parameters and provide a fair comparison between true IMC data and predicted IMC data. Full tissue IMC parameter predictions for the DFU and tonsil biopsy tissue section were performed using the same parameters as the tiled ROIs from the benchmark study above.

## Key Resources Table

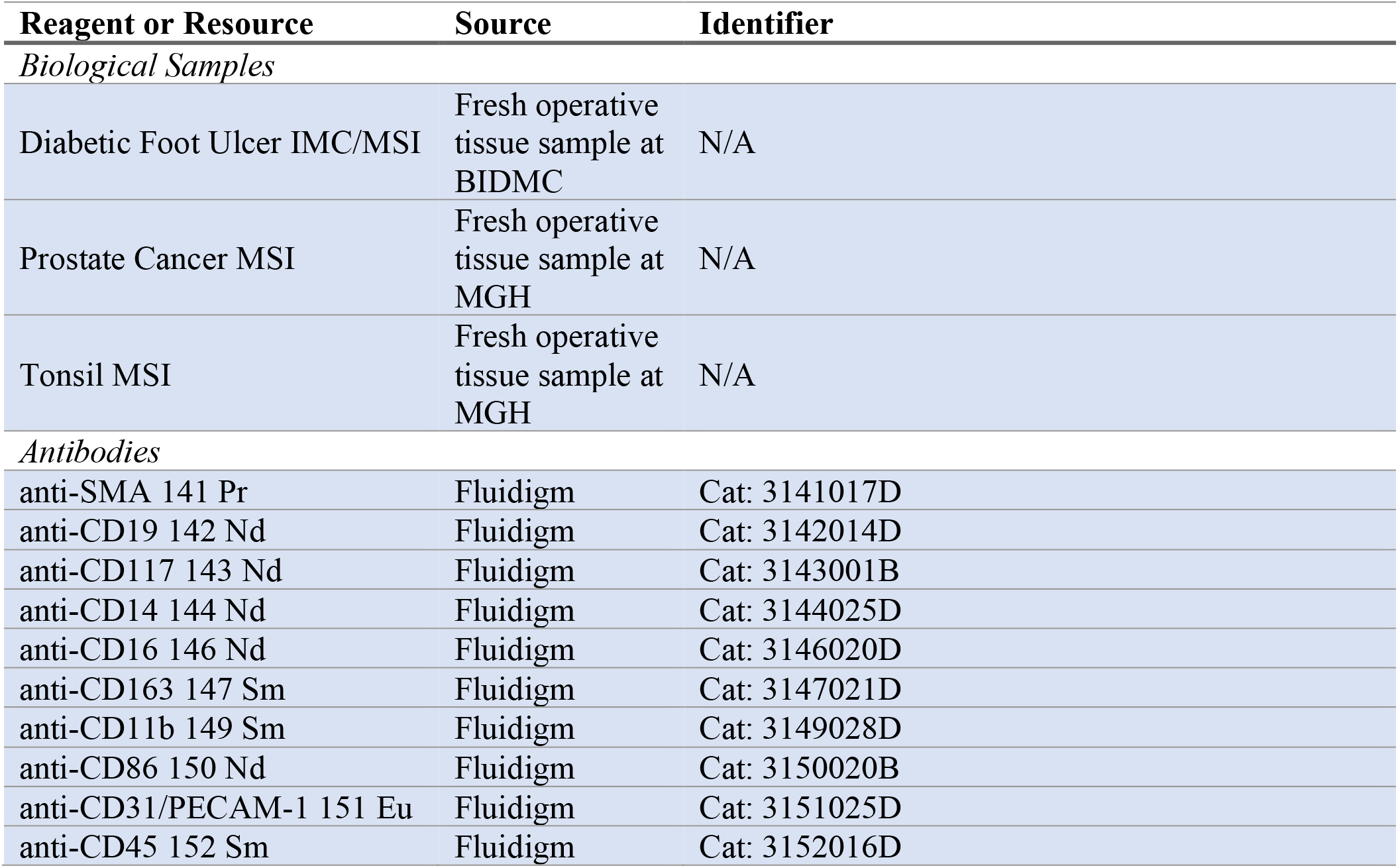
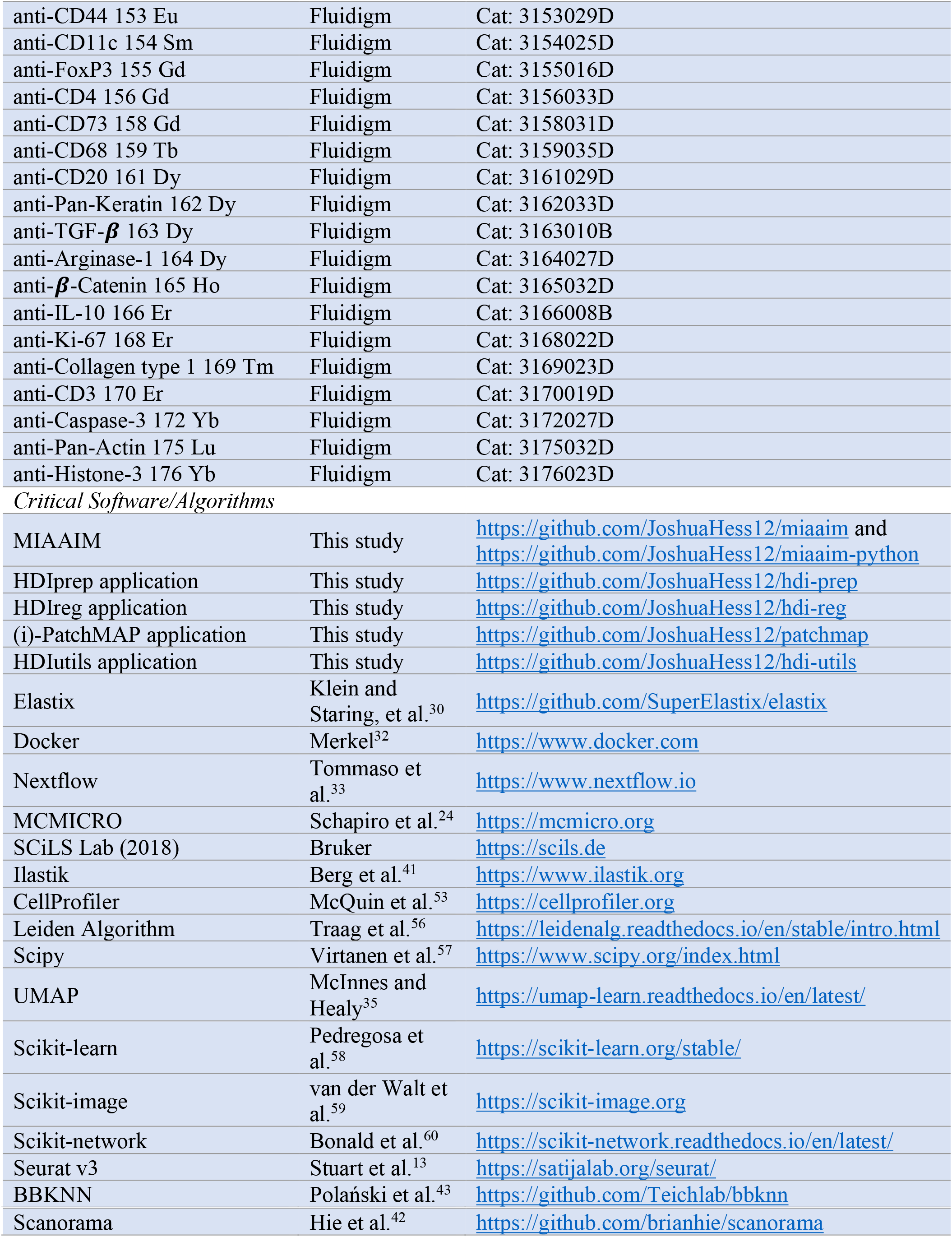

| Reagent or Resource | Source | Identifier |
| --- | --- | --- |
| *Biological Samples* |  |  |
| Diabetic Foot Ulcer IMC/MSI | Fresh operative tissue sample at BIDMC | N/A |
| Prostate Cancer MSI | Fresh operative tissue sample at MGH | N/A |
| Tonsil MSI | Fresh operative tissue sample at MGH | N/A |
| *Antibodies* |  |  |
| anti-SMA 141 Pr | Fluidigm | Cat: 3141017D |
| anti-CD19 142 Nd | Fluidigm | Cat: 3142014D |
| anti-CD117 143 Nd | Fluidigm | Cat: 3143001B |
| anti-CD14 144 Nd | Fluidigm | Cat: 3144025D |
| anti-CD16 146 Nd | Fluidigm | Cat: 3146020D |
| anti-CD163 147 Sm | Fluidigm | Cat: 3147021D |
| anti-CD11b 149 Sm | Fluidigm | Cat: 3149028D |
| anti-CD86 150 Nd | Fluidigm | Cat: 3150020B |
| anti-CD31/PECAM-1 151 Eu | Fluidigm | Cat: 3151025D |
| anti-CD45 152 Sm | Fluidigm | Cat: 3152016D |
| anti-CD44 153 Eu | Fluidigm | Cat: 3153029D |
| anti-CD11c 154 Sm | Fluidigm | Cat: 3154025D |
| anti-FoxP3 155 Gd | Fluidigm | Cat: 3155016D |
| anti-CD4 156 Gd | Fluidigm | Cat: 3156033D |
| anti-CD73 158 Gd | Fluidigm | Cat: 3158031D |
| anti-CD68 159 Tb | Fluidigm | Cat: 3159035D |
| anti-CD20 161 Dy | Fluidigm | Cat: 3161029D |
| anti-Pan-Keratin 162 Dy | Fluidigm | Cat: 3162033D |
| anti-TGF-β 163 Dy | Fluidigm | Cat: 3163010B |
| anti-Arginase-1 164 Dy | Fluidigm | Cat: 3164027D |
| anti-β-Catenin 165 Ho | Fluidigm | Cat: 3165032D |
| anti-IL-10 166 Er | Fluidigm | Cat: 3166008B |
| anti-Ki-67 168 Er | Fluidigm | Cat: 3168022D |
| anti-Collagen type 1 169 Tm | Fluidigm | Cat: 3169023D |
| anti-CD3 170 Er | Fluidigm | Cat: 3170019D |
| anti-Caspase-3 172 Yb | Fluidigm | Cat: 3172027D |
| anti-Pan-Actin 175 Lu | Fluidigm | Cat: 3175032D |
| anti-Histone-3 176 Yb | Fluidigm | Cat: 3176023D |
| *Critical Software/Algorithms* |  |  |
| MIAAIM | This study | https://github.com/JoshuaHess12/miaaim and https://github.com/JoshuaHess12/miaaim-python |
| HDIprep application | This study | https://github.com/JoshuaHess12/hdi-prep |
| HDIreg application | This study | https://github.com/JoshuaHess12/hdi-reg |
| (i)-PatchMAP application | This study | https://github.com/JoshuaHess12/patchmap |
| HDIutils application | This study | https://github.com/JoshuaHess12/hdi-utils |
| Elastix | Klein and Staring, et al.<sup>30</sup> | https://github.com/SuperElastix/elastix |
| Docker | Merkel<sup>32</sup> | https://www.docker.com |
| Nextflow | Tommaso et al.<sup>33</sup> | https://www.nextflow.io |
| MCMICRO | Schapiro et al.<sup>24</sup> | https://mcmicro.org |
| SCiLS Lab (2018) | Bruker | https://scils.de |
| Ilastik | Berg et al.<sup>41</sup> | https://www.ilastik.org |
| CellProfiler | McQuin et al.<sup>53</sup> | https://cellprofiler.org |
| Leiden Algorithm | Traag et al.<sup>56</sup> | https://leidenalg.readthedocs.io/en/stable/intro.html |
| Scipy | Virtanen et al.<sup>57</sup> | https://www.scipy.org/index.html |
| UMAP | McInnes and Healy<sup>35</sup> | https://umap-learn.readthedocs.io/en/latest/ |
| Scikit-learn | Pedregosa et al.<sup>58</sup> | https://scikit-learn.org/stable/ |
| Scikit-image | van der Walt et al.<sup>59</sup> | https://scikit-image.org |
| Scikit-network | Bonaldi et al.<sup>60</sup> | https://scikit-network.readthedocs.io/en/latest/ |
| Seurat v3 | Stuart et al.<sup>13</sup> | https://satijalab.org/seurat/ |
| BBKNN | Polański et al.<sup>43</sup> | https://github.com/Teichlab/bbknn |

## Data availability statement

All newly generated data from this study are available at https://www.synapse.org/MIAAIM_manuscript.

## Code availability statement

All software and code that produced the findings of the study, including all main and supplemental figures, are free and publicly available via GitHub at https://github.com/JoshuaHess12/miaaim-development. A user manual and training guide to get started using the MIAAIM software and associated workflows is available at https://joshuahess12.github.io/miaaim-python. Python modules that allow for individual applications of MIAAIM workflows can be found at https://github.com/JoshuaHess12/hdi-prep, https://github.com/JoshuaHess12/hdi-reg, and https://github.com/JoshuaHess12/patchmap. A Python package implementing MIAAIM is available at https://github.com/JoshuaHess12/miaaim-python.

## Author contribution statements

J.M.H., M.C.P., P.M.R. and R.F.S. designed the study. J.M.H. conceived and designed software and algorithms. R.F.S., P.M.R., and M.C.P. supervised the work. R.F.S. and P.M.R. conceived and performed MSI, IMC, and H&E multimodal imaging. J.M.H. performed processing and analysis of MSI, IMC, H&E, and CyCIF images. J.M.H., R.F.S., I.I., M.C.P. and P.M.R. designed benchmark studies and interpreted analyses. J.M.H. and D.S developed and implemented cell segmentation and quantification pipeline. J.M.H, J.J.I, and I.I. critically reviewed mathematics of the paper. M.S.R., G.T., C.L.W., A.V., N.Y.R.A. provided MSI, IMC, and H&E images. W.M.A. and A.E.S. provided critical input. J.M.H., I.I., M.C.P, P.M.R., and R.F.S. wrote the manuscript. J.M.H, I.I., M.C.P., P.M.R, and R.F.S. finalized the manuscript with input from all authors.

## ACKNOWLEDGEMENTS

J.M.H, M.C.P., P.M.R. and R.F.S. were supported by the Vaccine and Immunotherapy Center Innovation and Education Funds at Massachusetts General Hospital. J.M.H was previously supported and PMR is also supported by a Congressionally Directed Medical Research Program grant W81XWH-20-1-0301. J.M.H. is supported by a National Science Foundation Graduate Research Fellowship under grant no. 1746886. D.S. was supported by the University of Zurich BioEntrepreneur-Fellowship (BIOEF-17-001), a Swiss National Science Foundation Early Postdoc Mobility fellowship (P2ZHP3_181475, BMBF (01ZZ2004) and was a Damon Runyon Fellow supported by the Damon Runyon Cancer Research Foundation (DRQ-03-20). N.Y.R.A was partially supported by NIH grants U54-CA210180 and P41-EB028741. We thank Enterprise Research Infrastructure & Services at Partners HealthCare for their in-depth support and provision of computing resources.

## OUTSIDE INTERESTS

D.S. is a scientific consultant to Roche Glycart AG; none of these relationships are directly related to the topic of this study. R.F.S., J.M.H., P.M.R, and M.C.P. are authors on patent PCT/US21/48928 applied for by the General Hospital Corporation covering the described methods.

