## Supplementary Information for "MIAAIM: Multi-omics image integration and tissue state mapping using topological data analysis and cobordism learning"

### SUPPLEMENTARY FIGURES

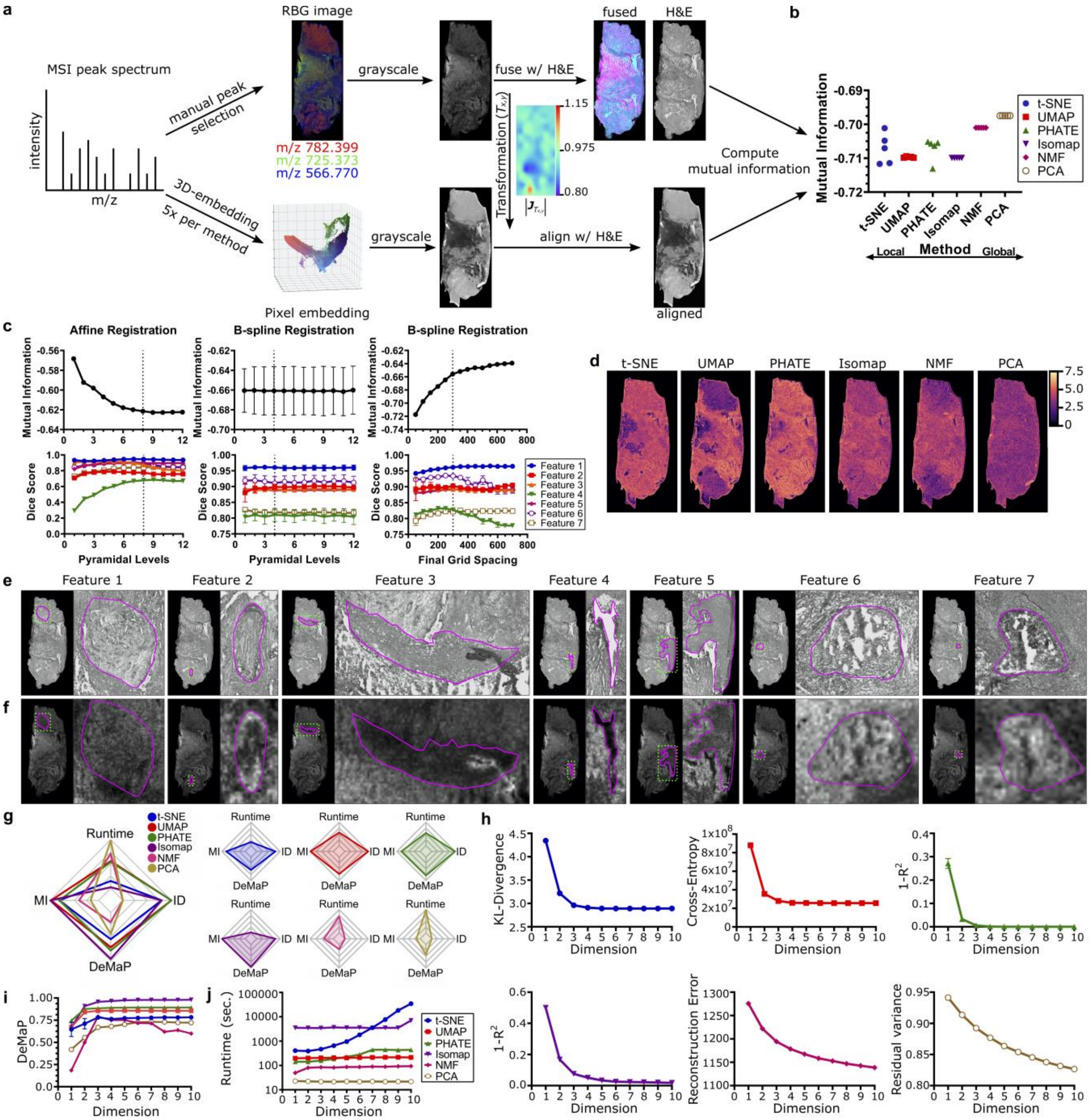


**Supplementary Fig. 1 | Performance of dimensionality reduction algorithms for summarizing diabetic foot ulcer mass spectrometry imaging data**

**a.** Three mass spectrometry peaks highlighting tissue morphology were manually chosen (top) and were used to create and RGB image representation of the MSI data, which was converted to a grayscale image.

The MSI grayscale image was then registered to its corresponding grayscale converted hematoxylin and eosin (H&E) stained section. The deformation field (middle), indicated by the determinant of its spatial Jacobian matrix, was saved to use downstream as a control registration. Three-dimensional Euclidean embeddings of the MSI data were then created using random initializations of each dimension reduction algorithm (bottom). These embeddings were then used to create an RGB image following the procedure above. The spatial transformation created by registering the manually identified peaks with the H&E image was then applied to dimension reduced grayscale images, aligning each to the grayscale H&E image. **b.** The mutual information of between each aligned grayscale embedded image (n = 5 per method) and the grayscale H&E image was calculated using Parzen window histogram density estimation with a histogram bin width of 64. Plot is oriented so that results are consistent with the notion of a "cost function" in optimization contexts, where the goal is to minimize cost. Thus, larger negative values depict higher mutual information. UMAP consistently captures multi-modal information content with respect to the H&E data. **c.** Optimization of image registration between the grayscale version of manually identified mass spectrometry peaks and the grayscale H&E image (**a**, top) using mutual information as a cost function with external validation using dice scores on 7 manually annotated regions. Registration parameters used for the final registration used in **a** are indicated with dashed lines. Registration was performed by first aligning images with a multi-resolution affine registration (left). The transformed grayscale version of manually identified mass spectrometry peaks was then registered to the grayscale H&E image using a nonlinear, multi-resolution registration. **d.** Average neighborhood entropy (n = 5) of each pixel calculated within a 10-pixel disc across dimension reduction algorithms. Results show UMAP's ability to highlight structure in the tissue section. **e.** Manual annotations of grayscale H&E image used for validating registration quality with controlled deformation field in **a** used for mutual information calculations in **b. f.** Cropped regions using the same spatial coordinates as **e** of manually annotated regions used to calculate the dice scores in **c**. Results show good spatial overlap across disparate annotations**. g.** Radar plots showing performance comparison of dimension reduction algorithms spanning a range of data representation – linear, nonlinear, local, and global data structure preservation (t-SNE, UMAP, PHATE, Isomap, NMF, PCA). Shown are mean values (n = 5) of algorithm runtime (top, log transformed), estimated steady-state manifold embedding dimensionality (right), noise robustness (bottom), and multi-modal mutual information to DFU MSI data (left). All plots are oriented so that larger values depict better algorithmic performance. Results show UMAP's ability to efficiently capture data complexity with few degrees of freedom while balancing noise robustness with multi-modal information content contained in histology images. **h.** Intrinsic dimensionality of MSI data estimated by each dimension reduction method. Embedding errors (y-axes) are not comparable across plots. Plotted are the mean and standard deviation (n = 5) embedding errors across embedding dimensions 1-10. Convergence on y-axes indicates that increasing the dimension of the resulting embeddings no longer improves an algorithms ability to capture data complexity. Results show that the intrinsic dimensionality estimated by nonlinear methods (t-SNE, UMAP, PHATE, Isomap) is far less than that of linear methods (NMF, PCA), meaning that fewer dimensions are needed to accurately describe the data set. **i.** Denoised manifold preservation (DeMaP) metric between Euclidean distances in resulting embeddings corresponding to non-peak-picked data and geodesic distances in ambient space (not dimension reduced after peak-picking) of corresponding peak-picked data. Results showing the mean and standard deviation DeMaP metric (Spearman's rho correlation coefficient) for all tested dimension reduction methods (n = 5). Nonlinear methods Isomap, PHATE, and UMAP all consistently preserve manifold structure without prior filtering of the data with consistent correlations greater than 0.85 across dimensions 2-10. **j.** Computational runtime for each algorithm across embedding dimensions 1-10. Plotted are the mean and standard deviation (n = 5) across each number of dimensions for each method. Nonlinear methods t-SNE and Isomap require longer run times than the nonlinear methods PHATE and UMAP. Linear methods require the least amount of run time; however, they fail to capture data complexity succinctly.

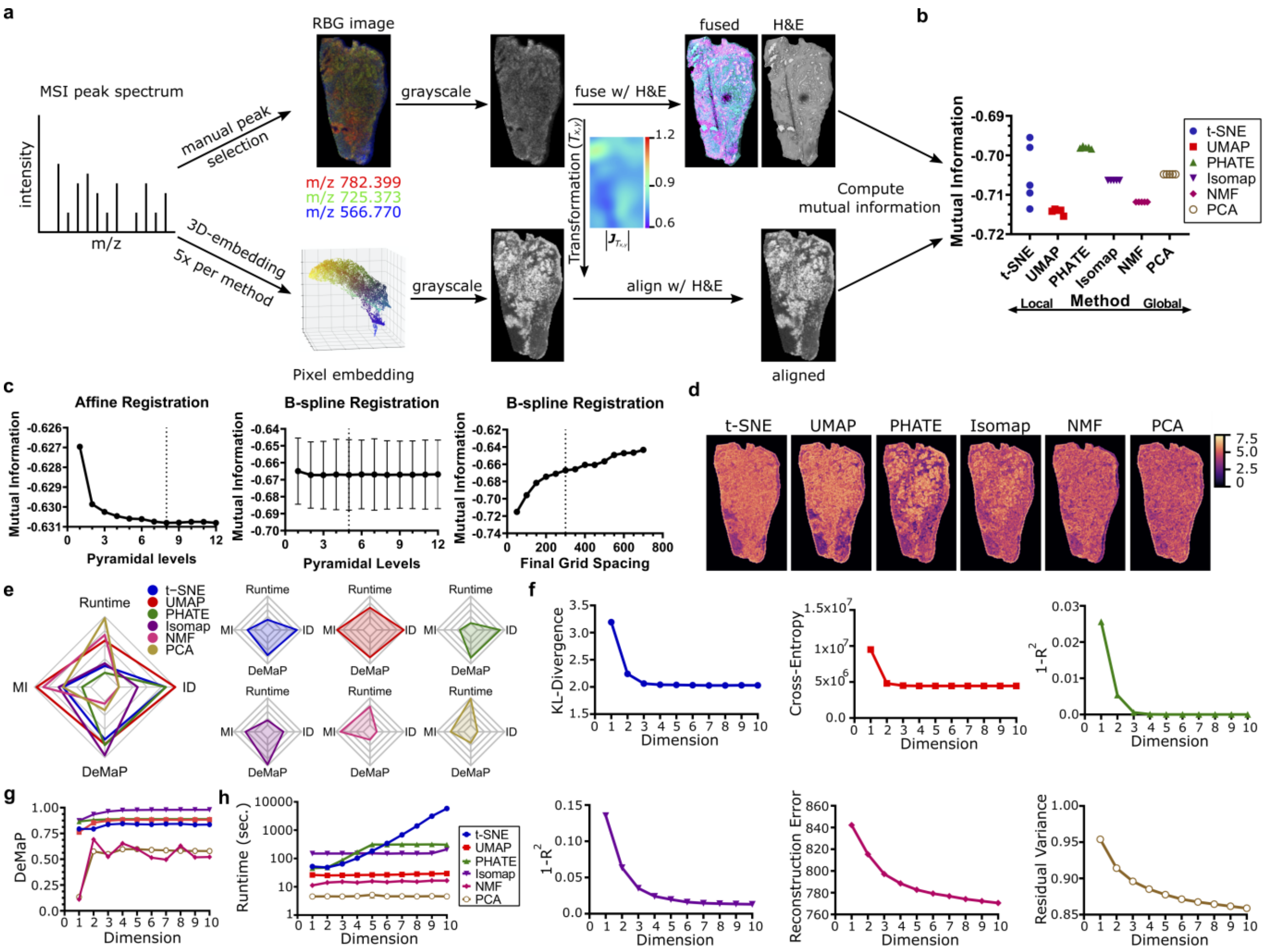


**Supplementary Fig. 2 | Performance of dimensionality reduction algorithms for summarizing prostate cancer mass spectrometry imaging data**

**a.** Same as **Supplementary Fig. 1a** for prostate cancer tissue biopsy. **b.** Same as **Supplementary Fig. 1b** for prostate cancer tissue biopsy. **c.** Optimization of image registration between the grayscale version of manually identified mass spectrometry peaks and the grayscale H&E image (**a**, top) using mutual information as a cost function. Registration parameters used for the final registration used in **a** are indicated with dashed lines. Registration was performed by first aligning images with a multi-resolution affine registration (left). The transformed grayscale version of manually identified mass spectrometry peaks was then registered to the grayscale H&E image using a nonlinear, multi-resolution registration. **d.** Same as **Supplementary Fig. 1d** for prostate cancer tissue biopsy. **e.** Same as **Supplementary Fig. 1g** for prostate cancer tissue biopsy. **f.** Same as **Supplementary Fig. 1h** for prostate cancer tissue biopsy. **g.** Same as **Supplementary Fig. 1i** for prostate cancer tissue biopsy. Nonlinear methods Isomap, PHATE, and UMAP all consistently preserve manifold structure without prior filtering of the data with consistent correlations greater than 0.75 across dimensions 2-10. **h.** Same as **Supplementary Fig. 1j** for prostate cancer tissue biopsy

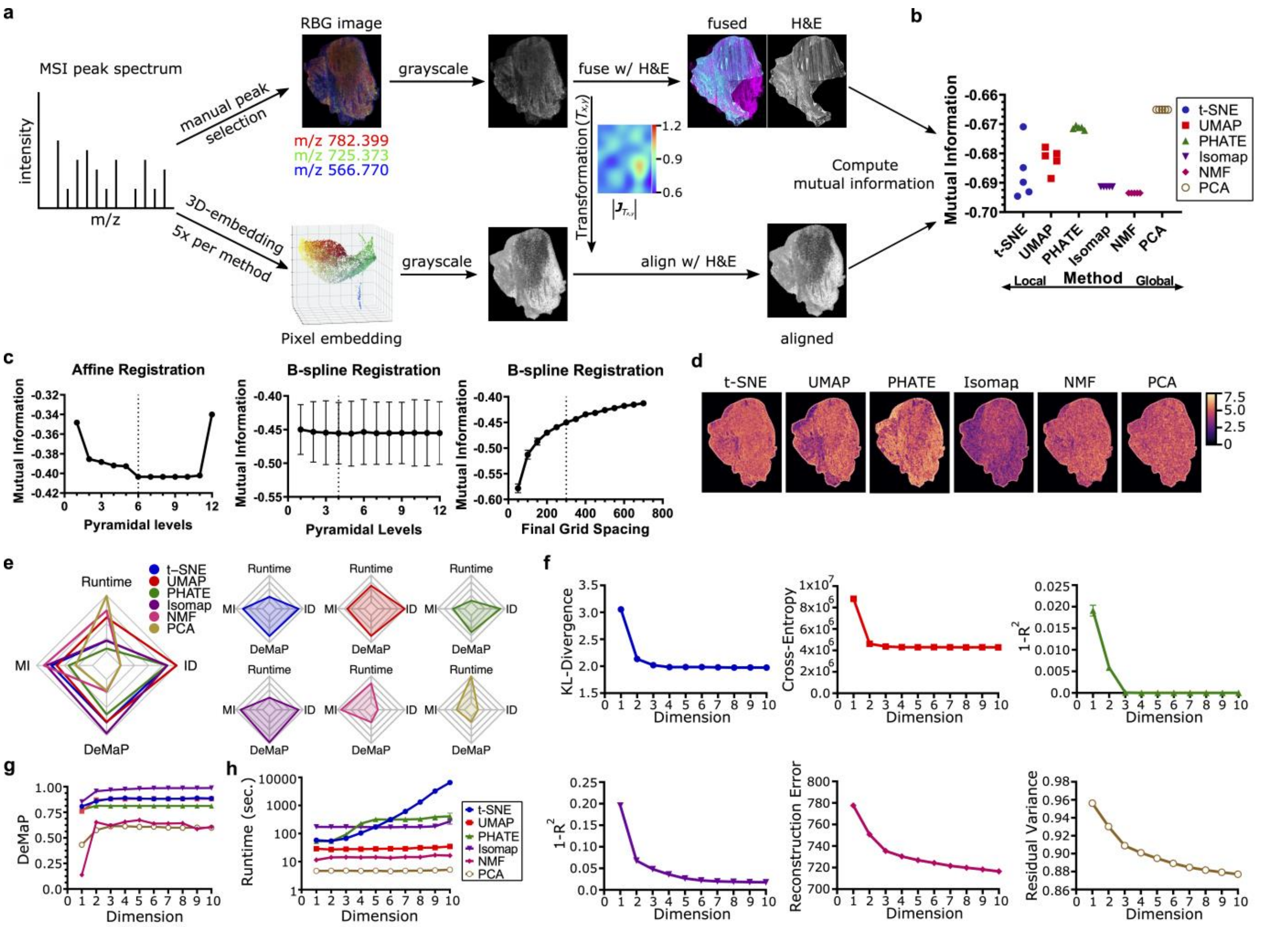


**Supplementary Fig. 3 | Performance of dimensionality reduction algorithms for summarizing tonsil mass spectrometry imaging data**

**a.** Same as **Supplementary Fig. 1a** for tonsil tissue biopsy. **b.** Same as **Supplementary Fig. 1b** for tonsil tissue biopsy. Isomap and NMF consistently capture multi-modal information content with respect to the H&E data. **c.** Same as **Supplementary Fig. 2c** for tonsil tissue biopsy. **d.** Same as **Supplementary Fig. 1d** for tonsil tissue biopsy. **e.** Same as **Supplementary Fig. 1g** for tonsil tissue biopsy. **f.** Same as **Supplementary Fig. 1h** for tonsil tissue biopsy. **g.** Same as **Supplementary Fig. 1i** for tonsil tissue biopsy. **h.** Same as **Supplementary Fig. 1j** for tonsil tissue biopsy.

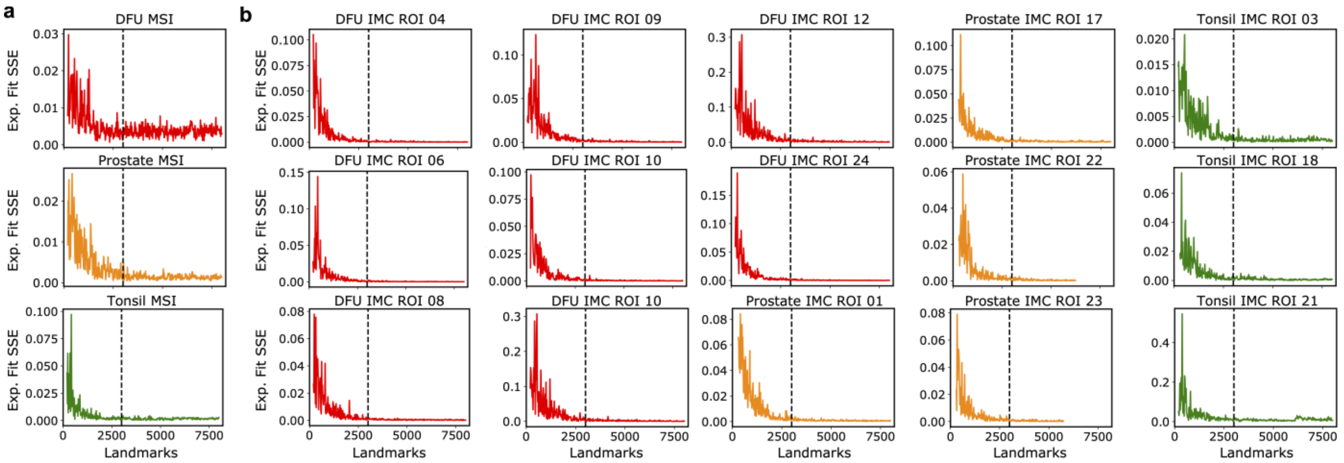


**Supplementary Fig. 4 | Spectral centroid landmarks recapitulate steady-state manifold embedding dimensionalities across tissue types and imaging technologies**

**a.** Sum of squared errors of exponential regressions fit to steady state embedding dimensionality selections from spectral landmarks compared to full mass spectrometry imaging data sets across DFU, Prostate, and Tonsil tissues. Discrepancies between exponential regressions fit to the cross-entropy of landmark centroid embeddings and full data set embeddings approach zero as the number of landmarks increases. Dashed lines show MIAAIM’s default selection of 3,000 landmarks for computing steady-state manifolds embedding dimensionalities. **b**. Same as **a** for subsampled pixels in imaging mass cytometry regions of interest.

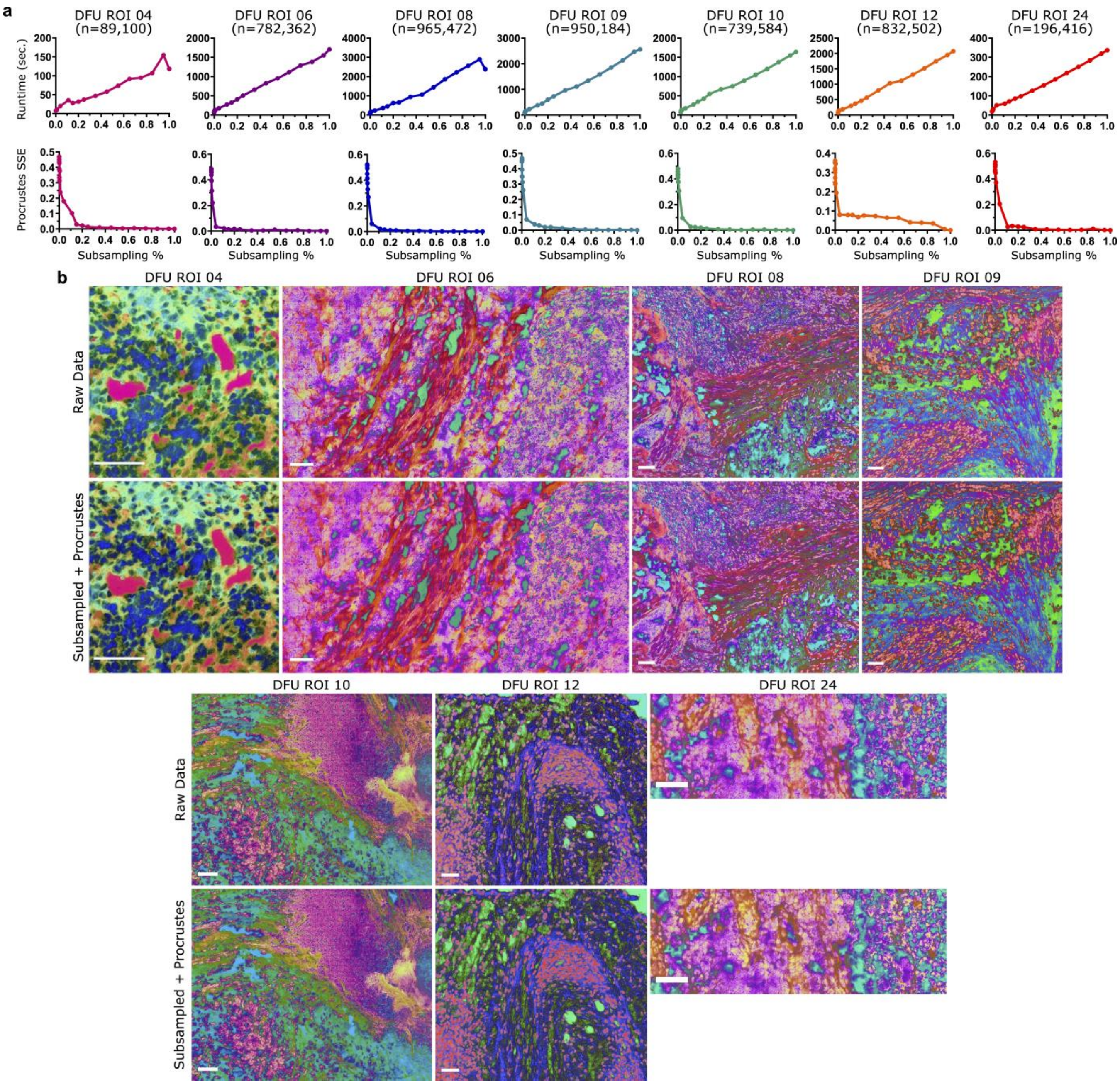


**Supplementary Fig. 5 | UMAP embeddings of spatially subsampled imaging mass cytometry data with out-of-sample projection recapitulate full data embeddings while decreasing runtime in DFU samples**

**a.** Three-dimensional UMAP embedding (HDIprep compression) runtime plotted with respect to subsampling percentages of IMC ROIs from the DFU tissue biopsy (top). Procrustes transformation sum of squared errors after transforming subsampled embedding to the full pixel embedding across subsampling percentages (bottom). **b.** Comparison of RGB (red, green, blue) images created by reconstructing images from pixel embeddings on all data (top) versus subsampled data with subsequent out-of-sample projection and Procrustes transformation to align subsampled embedding to the full pixel embedding (bottom) (scale bars = 80 $\mu$m). Subsampling percentages of images shown in bottom row of **b** correspond to MIAAIM default parameters that based on number of pixels in images.

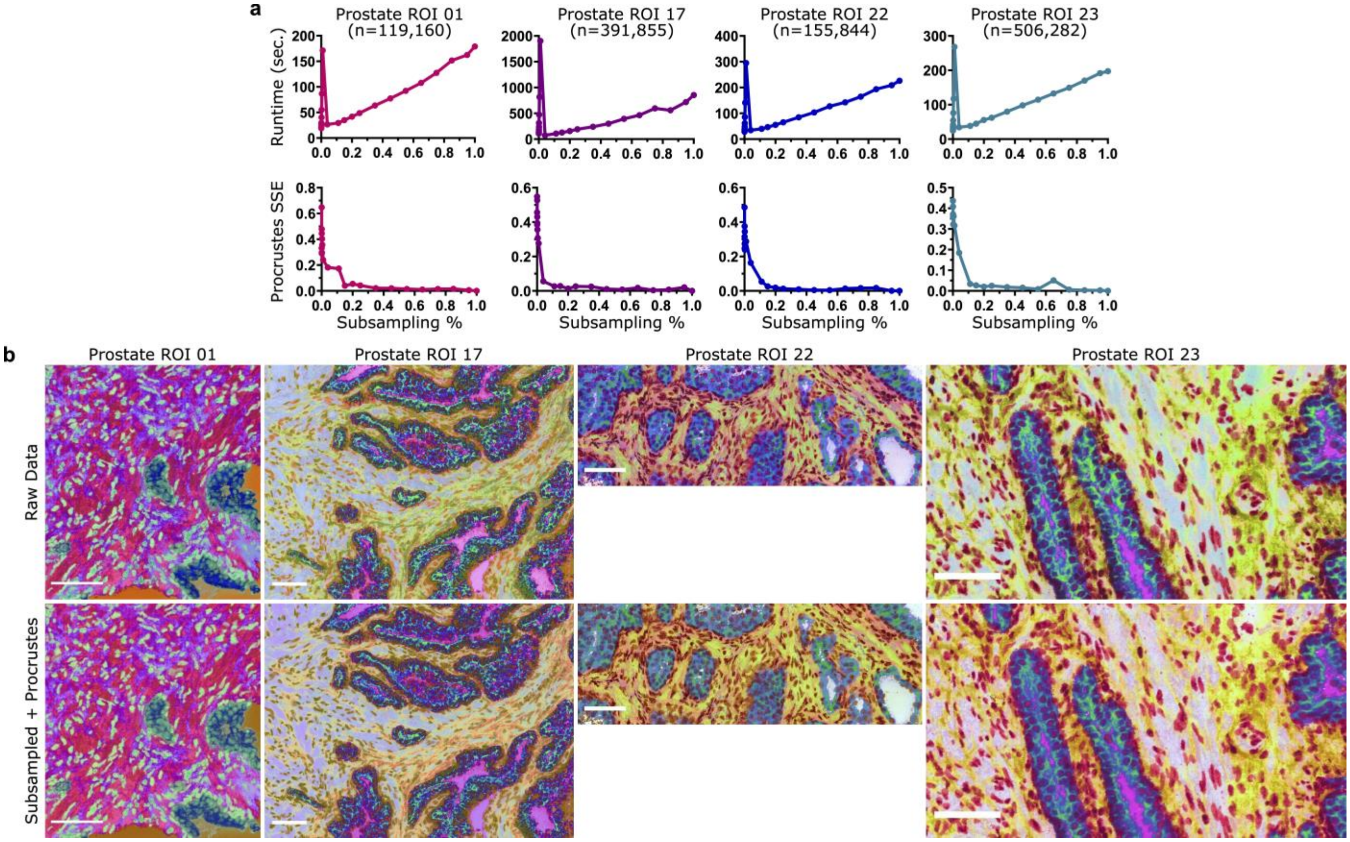


**Supplementary Fig. 6 | UMAP embeddings of spatially subsampled imaging mass cytometry data with out-of-sample projection recapitulate full data embeddings while decreasing runtime in prostate cancer samples**

**a-b.** Same as **Supplementary Fig. 5** for prostate tumor tissue biopsy IMC ROIs.

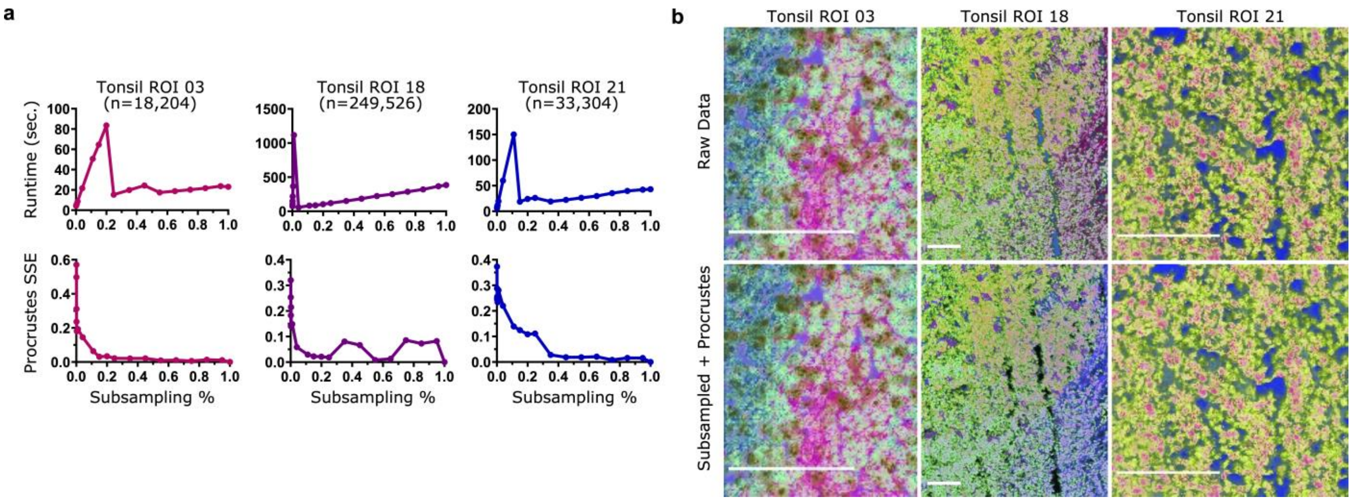


**Supplementary Fig. 7 | UMAP embeddings of spatially subsampled imaging mass cytometry data with out-of-sample projection recapitulate full data embeddings while decreasing runtime in tonsil samples**

**a-b.** Same as **Supplementary Fig. 5** for tonsil tissue biopsy IMC ROIs.

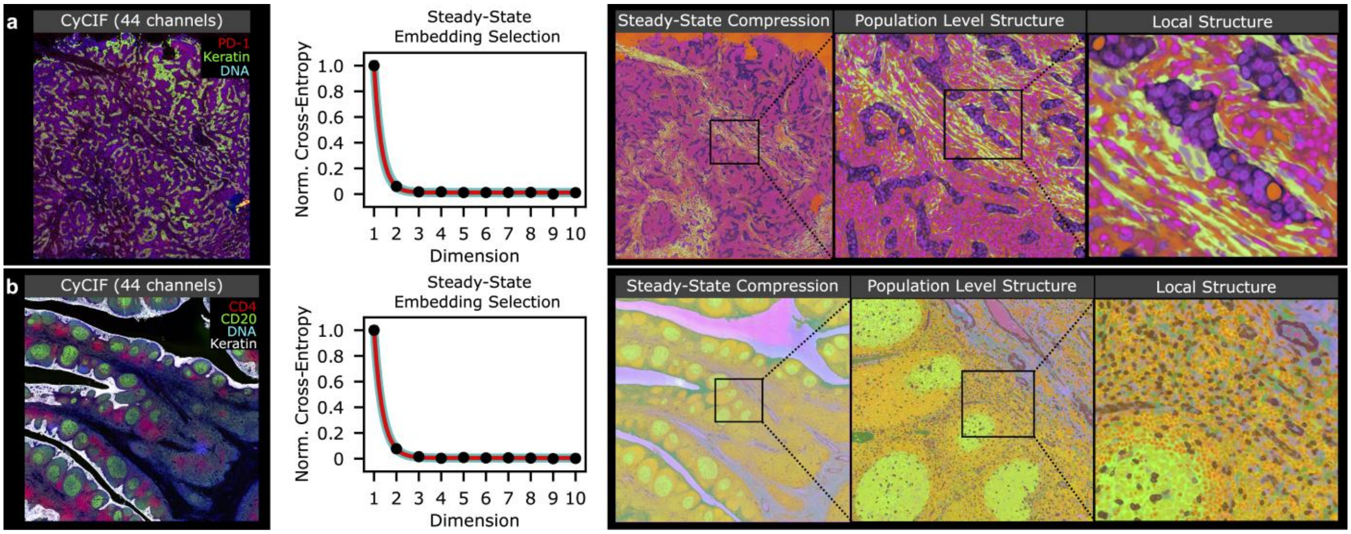


**Supplementary Fig. 8 | MIAAIM image compression scales to large fields of view and high-resolution multiplexed image datasets by incorporating parametric UMAP**

**a.** Multiplex CyCIF image of lung adenocarcinoma metastasis to the lymph node (n = ~100 million pixels, 0.65 $\mu$m/pixel resolution, 44 channels, 27 antibodies) and corresponding steady-state UMAP embedding and spatial reconstruction (shown are three UMAP channels of 4 channel steady state embedding). Parametric UMAP compresses millions of pixels and preserves tissue structure across multiple length scales. **b.** Same as **Supplementary Fig. 8a** for tonsil CyCIF data (n = ~256 million pixels, 0.65 $\mu$m/pixel resolution).

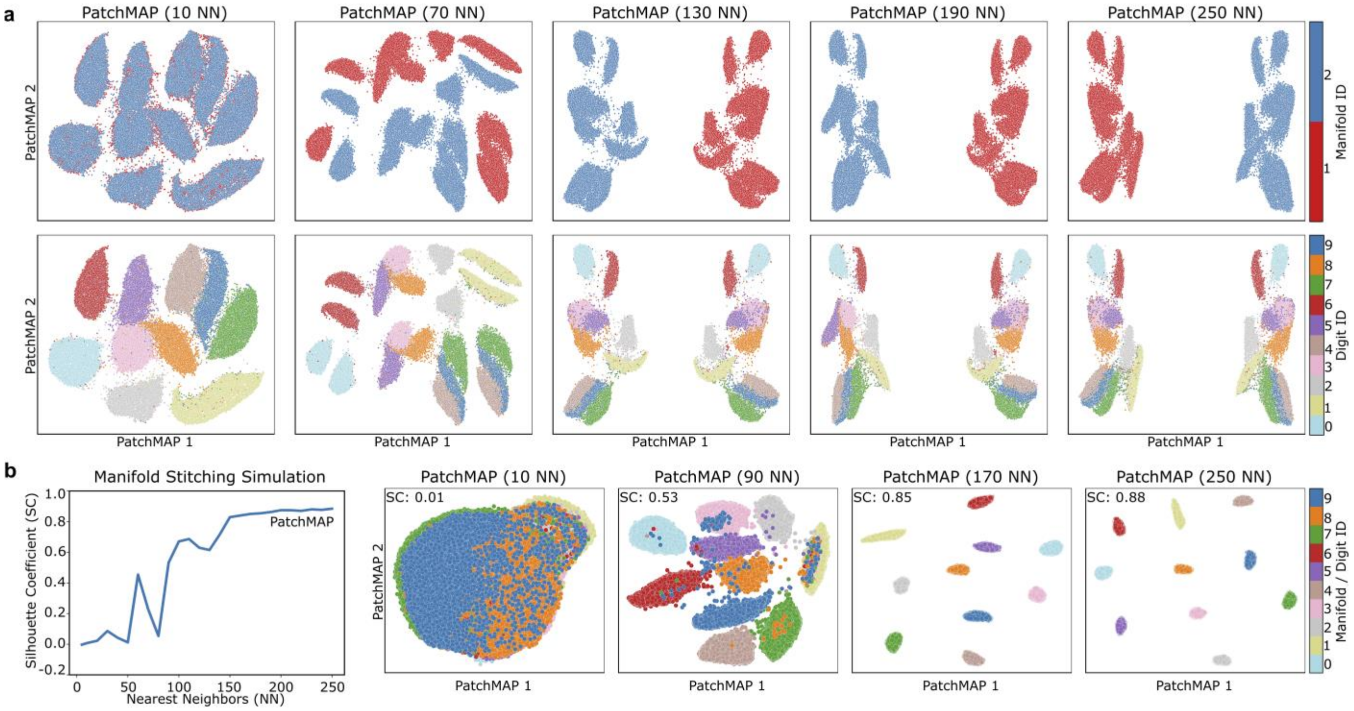


**Supplementary Fig. 9 | PatchMAP preserves boundary manifold structure while accurately embedding inter-boundary manifold relationships in cobordisms**

**a.** PatchMAP embedding of MNIST digits dataset (n = 70,000) randomly split into two equally-sized boundary manifolds. At lower values of nearest neighbors, PatchMAP resembles UMAP embeddings because open neighborhoods are preserved after intersecting pairwise nearest neighbor queries. Under these conditions, the intersection operation resembles the fuzzy set union that UMAP implements. At higher values of nearest neighbors, PatchMAP captures manifold relationships in cobordisms while preserving boundary manifold structure. Here, PatchMAP aligns boundary manifolds along a primary axis to produce near mirror images. This reflects the equal splitting of the data in half, and it is captured in cobordism geodesic distances. **b.** Validation of **Fig. 4b** on the full MNIST digits dataset, where each digit in the dataset is considered to be a boundary manifold. Lower values of nearest neighbors resemble UMAP embeddings, and higher values of nearest neighbors allow PatchMAP to accurately model cobordism geodesic distances.

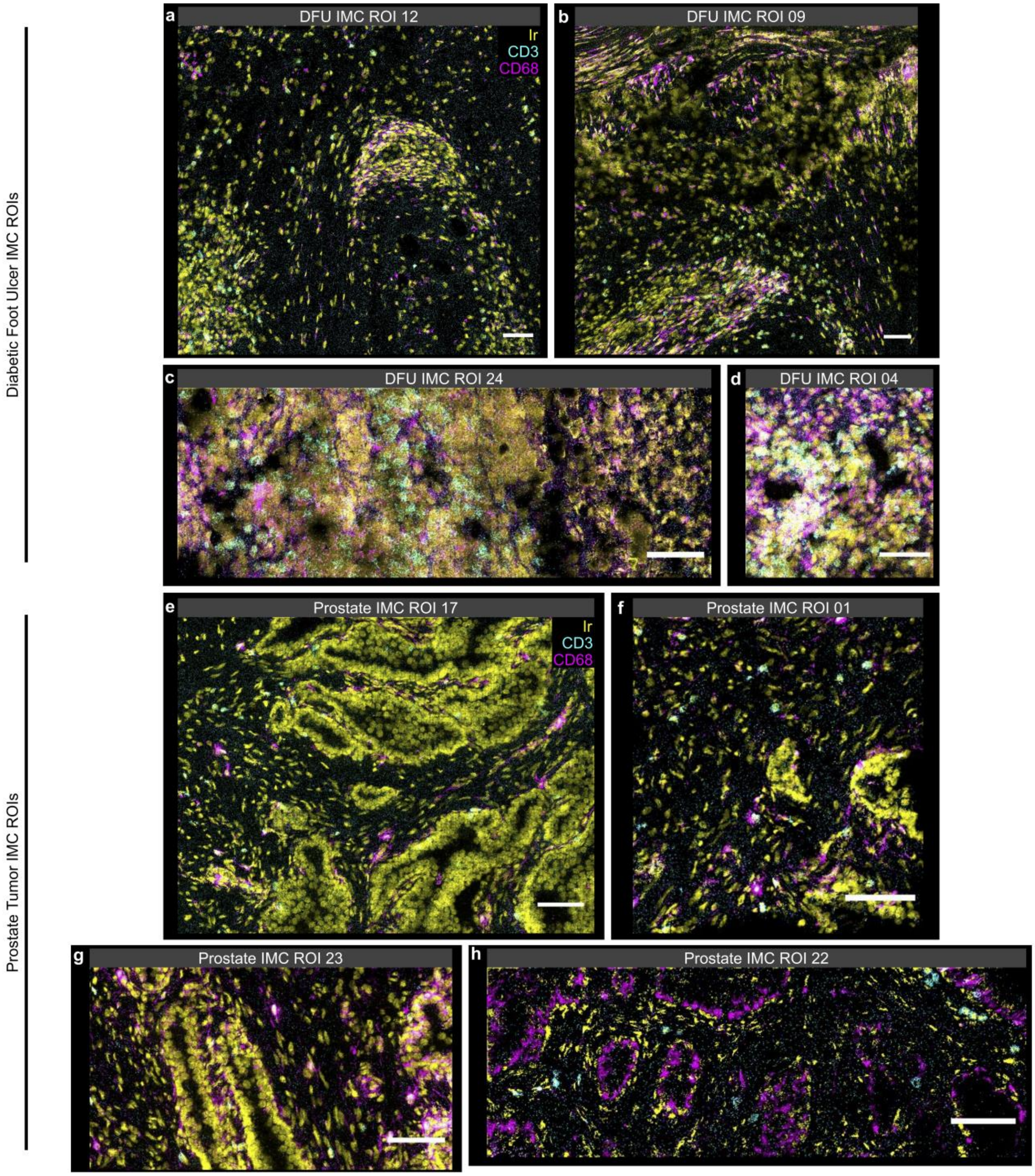


**Supplementary Fig. 10 | Sparse immune cell infiltration in prostate tumor biopsy IMC ROIs supports lack of boundary manifold connections between DFU and prostate**

**a-d.** DFU tissue biopsy IMC ROIs farthest from the center point of the ulcer that exhibit dense areas of immune cell populations (yellow, Iridium; cyan, CD3; magenta, CD68; scale bars = 80 $\mu$m). **e-h.** Prostate tumor tissue biopsy IMC ROIs showing sparse immune cell infiltration (yellow, Iridium; cyan, CD3; magenta, CD68; scale bars = 80 $\mu$m).

* Contrast and intensity of channels in each image individually adjusted for visualization purposes.

#### SUPPLEMENTARY TABLES

| Imaging Technology | Input File Format | Parameters Detected | Tested with MIAAIM | Type of Processing | Data Set Reference |
|---|---|---|---|---|---|
| **MSI**[1] | imzML | Proteins, Lipids, Metabolites | ✓ | Compression | This study |
| **Histological Stains (e.g. H&E)** | TIF(F) | Tissue morphology | ✓ | Denoising | This study |
| **IMC**[2] | OME-TIF(F) | Subcellular proteomics | ✓ | Compression | This study |
| **MIBI**[3] | TIF(F) | Subcellular proteomics | ✓ | Compression | 4 |
| **CODEX**[5] | TIF(F) Hyper Stack | Subcellular proteomics | ✓ | Compression | 5 |
| **CyCIF**[6] | OME-TIF(F) | Subcellular proteomics | ✓ | Compression | 7 |
| **4i**[8] | - | Sub-organelle proteomics | ✗ | Compression | - |
| **Slide-seq**[9] | - | Subcellular Transcriptomics | ✗ | Compression | - |
| **RNA-ISH related technologies** | - | Transcriptomics | ✗ | Compression | - |

**Supplementary Table 1 | Current imaging technologies anticipated to be compatible and those tested for compatibility with MIAAIM image compression and pre-processing for subsequent alignment**

#### Supplementary Table References


1. McDonnell, L.A. & Heeren, R.M. Imaging mass spectrometry. *Mass spectrometry reviews* **26**, 606-643 (2007).
2. Giesen, C. et al. Highly multiplexed imaging of tumor tissues with subcellular resolution by mass cytometry. *Nature methods* **11**, 417-422 (2014).
3. Angelo, M. et al. Multiplexed ion beam imaging of human breast tumors. *Nature medicine* **20**, 436 (2014).
4. https://github.com/ionpath/mibilib.
5. Goltsev, Y. et al. Deep profiling of mouse splenic architecture with CODEX multiplexed imaging. *Cell* **174**, 968-981. e915 (2018).
6. Lin, J.-R. et al. Highly multiplexed immunofluorescence imaging of human tissues and tumors using t-CyCIF and conventional optical microscopes. *Elife* **7** (2018).
7. Rashid, R. et al. Highly multiplexed immunofluorescence images and single-cell data of immune markers in tonsil and lung cancer. *Scientific data* **6**, 1-10 (2019).
8. Gut, G., Herrmann, M.D. & Pelkmans, L. Multiplexed protein maps link subcellular organization to cellular states. *Science* **361** (2018).
9. Rodriques, S.G. et al. Slide-seq: A scalable technology for measuring genome-wide expression at high spatial resolution. *Science* **363**, 1463-1467 (2019).

### SUPPLEMENTARY NOTES

#### Supplementary Note 1

#### Combining MIAAIM with existing bioimaging analysis software

MIAAIM's core functionality enables cross-technology and cross-tissue comparisons. As shown in our MSI and IMC multi-modal imaging integration examples, it has broad applications that may be configured and executed using other software applications. We anticipate that a challenge for many users will be the execution of sequential registration and analysis across varied software. MIAAIM's multiple output data formats interface with a number of tools for visualization, cell segmentation, and single-cell analysis, creating an avenue for continued investigation of multimodal tissue portraits in various settings. Examples of compatible software include MCMICRO[10] (BioFormats input), histoCAT[11] (TIFF input), Ilastik[12] (HDF5 input), ImageJ/FIJI[13] (BioFormats and NIfTI input), Qupath[14] (BioFormats input), and napari[15] (image visualization software).

#### Supplementary Note 2

#### Notes on the HDIreg workflow's expected performance

A fundamental assumption in intensity-based image registration is that quantifiable relationships exist between modalities – this is often met in practice, as shown in our MSI and IMC multi-modal imaging data applications. However, this assumption can be compromised by artefacts, such as folds, tears, and, in the case of serial sectioning, nonlinear deformations. In our experience, glandular tissues, such as those derived from prostate, likely show high structural variability over short distances, making the alignment of images from distinct sections challenging. Manual landmark guidance can be used in difficult use-cases such as those posed by serial tissue sectioning. By using the Elastix library, HDIreg also offers a multitude of similarity measures for single-channel registration, in addition to the manifold alignment scheme used for multichannel registration. We note that histogram-based mutual information has outperformed KNN $\alpha$-MI in these single-channel registration settings[16], which we used in our benchmark studies.

#### Supplementary Note 3

#### HDIprep dimension reduction validation

**Dimensionality reduction algorithm benchmarking (Supplementary Figs. 1-3).**

Our investigation involved a range of dimension reduction methods, spanning from local nonlinear methods, to global, linear methods. Considered methods included t-distributed stochastic neighbor embedding (t-SNE)[17], uniform manifold approximation and projection (UMAP)[18], potential of heat diffusion for affinity-based transition embedding (PHATE)[19], isometric mapping (Isomap)[20], non-negative matrix factorization (NMF)[21], and principle components analysis (PCA)[22].

To assess each methods' ability to provide an appropriate data representation while enabling multi-modal correspondences, we measured their ability to (i.) generalize to arbitrary numbers of features or necessary degrees of freedom to accurately represent data modalities (ii.) succinctly capture data complexity (iii.) maximize the information content shared between imaging modalities (iv.) be robust to noise and (iv.) be computationally efficient.

**(i.-ii.) Estimating intrinsic data dimensionality.** To identify an appropriate method for reducing the complexity of the mass-spectrometry based image data sets, we hypothesized that introducing more degrees of freedom in the coordinates of embedded data (i.e. increases in the dimension of embeddings) would result in increases of the similarity between each methods' embedding and its high-dimensional counterpart with respect to the objective function of each algorithm. Therefore, we viewed each algorithm separately with a distinct objective function, and identified the appropriate target dimensionality for the data to be embedded in for each method by analyzing the objective function errors produced by each method after embedding the data in increasing dimensions. To do this, we created a suitable score for estimating the error associated with embedding the MSI data in Euclidean $n$-space, $\mathbb{R}^n$, for each dimension reduction method across tissue types and ascending embedding dimensions. For this analysis, we focused on the MSI data rather than the IMC data, which we found was not feasible to apply most dimension reduction methods to because of data size (number of pixels/high resolution).

To determine each method's estimated intrinsic dimensionality of the data set, we identified the point in each methods' error graph where increases in dimensionality no longer reduced embedding error. To do this, we viewed increases in the dimensionality of real-valued data in a natural way by modelling increases in dimensionality as exponential increases in potential positions of points (i.e., increasing copies of the real line, $\mathbb{R}^n$). We therefore fit a least-squares exponential regression to the error curves of data embedding, and 95% confidence intervals (CI) were constructed by modelling gaussian residual processes. The optimal embedding dimensions for each method were selected by simulating samples along the expected value of the fit curve and identifying the first integer-valued instance that fell within the 95%

CI for the exponential asymptote. In this way, the minimum degrees of freedom necessary to capture data complexity was identified. The average error curves for each method across 5 random initializations of each algorithm across each MSI data set are shown in **Supplementary Figs. 1, 2, and 3.** The methods and rationale used for calculating each method's embedding error are outlined below:

UMAP. The UMAP algorithm falls in the category of manifold learning techniques, and it aims to optimize the embedding of a fuzzy simplicial set representation of high-dimensional data into lower dimensional Euclidean spaces. Practically, a low dimensional fuzzy simplicial set is optimized so that the fuzzy set cross-entropy between its high-dimensional counterpart is minimized. The fuzzy-set cross entropy is defined explicitly in **Definition 1, Methods**, given by McInnes and Healy[18].

While the theoretical underpinnings of UMAP are grounded in category theory, the practical implementation of UMAP boils down to weighted graphs. To provide an estimate of the intrinsic dimensionality of the data determined by UMAP, we used the open-source implementation in Python[23] with 15 nearest neighbors, a value of 0.1 for minimum distance in the resulting embedding, and we allow the algorithm to optimize the embedding for the default value of 200 iterations for each dimension. The cross-entropy for each dimension between the high dimensional fuzzy simplicial set and the low dimensional counterpart was computed using a Python-converted module of the MATLAB UMAP implementation[24].

T-SNE. T-SNE is a manifold-based dimension reduction method that aims to preserve local structure in data sets for visualization purposes[17]. To achieve this, t-SNE minimizes the difference between distributions representing the local similarity between points in the original, high-dimensional ambient space and the respective low dimensional embedding. The difference between these two distributions is determined by the Kullback-Leibler (KL) divergence between them. As a result, we report the final value of the KL-divergence upon embedding as a means of estimating the error associated with t-SNE embeddings in each dimension. For all t-SNE calculations, we use an open-source multi-core implementation[25] with the default parameters (perplexity of 30).

Isomap. Isomap is a manifold-based dimension reduction method that uses classic multidimensional scaling (MDS) to preserve interpoint geodesic distances[20]. To do this, the geodesic distance between points are determined by shortest-path graph distances using the Euclidean metric. The pairwise distance matrix represented by this graph is then embedded into $n$-dimensional Euclidean space via classical MDS, a metric-preserving technique that finds the optimal transformation for inter-point Euclidean metric preservation. As a result of the implicit linearity in classic MDS, we estimate the intrinsic dimensionality of the data by calculating the reconstruction error in each dimension using $1 - R^2$, where $R$ is the standard linear correlation coefficient between the geodesic distance matrix and the pairwise Euclidean distance matrix in $\mathbb{R}^n$. For all calculations, 15 nearest neighbors were chosen for determining shortest-path graph distances, and the Minkowski metric with an order of two for the norm of the difference $||u - v||_p$ was chosen. All Isomap calculations were performed using Scikit-learn[26].

_PHATE._ PHATE is a manifold-based dimension reduction technique developed for data visualization that captures both global and local features of data sets[19]. PHATE achieves this by modelling relationships between data points as t-step random walk diffusion probabilities and by subsequently calculating potential distances between data points through comparison of each pair of points' respective diffusion distributions to all others in the data set[19]. These potential distances are then embedded in $n$-dimensional space using classic MDS followed by metric MDS. Metric MDS is suitable for embedding points with dissimilarities given by any metric, relaxing Euclidean constraints imposed by classical MDS, through minimizing the following stress function[19] $S$:

$$S(\hat{x}_1 \dots \hat{x}_N) = \sqrt{\frac{\sum_{i,j}\left(D_{x_i,x_j} - ||\hat{x}_i - \hat{x}_j||\right)^2}{\sum_{i,j}\left(D_{x_i,x_j}\right)^2}}$$

where $D$ is the metric defined over points $x_1 \dots x_N$ in the original data set, and $\hat{x}_1 \dots \hat{x}_N \epsilon \mathbb{R}^n$ are the corresponding embedded data points in dimension $n$. This stress function amounts to a least-squares optimization problem. In the scalable form of PHATE used for large data sets, landmarks instead of points are embedded in $n$-dimensional Euclidean space based on their pairwise potential distances using the above stress function. Out-of-sample embedding for all data points is performed by calculating linear combinations of the t-step transition matrix from points to landmarks using the embedded landmark coordinates as weights. If the stress function for metric MDS is zero, then the dimension reduction process is fully able to embed and capture the interpoint distances of the data. This would provide an error estimate to be used for analyses on intrinsic data dimension for the full data set and full PHATE algorithm; however, for the landmark-based calculations, not all points are embedded using metric MDS. Given the linear interpolation scheme and the initialization of scalable PHATE using classical MDS on landmark potential distances, we posited that the reconstruction error given by $1 - R^2$, where $R$ is the linear correlation coefficient between the point-to-landmark transition matrix and the pairwise Euclidean distance matrix in $\mathbb{R}^n$, provides an estimate for the error associated with embedding the full data set. All PHATE calculations were performed in Python using 15 nearest neighbors and the default number of 2,000 landmark points.

NMF. Non-negative matrix factorization (NMF)[21] is a linear dimension reduction technique that aims to minimize the divergence between an input matrix $\boldsymbol{X}$ and its reconstruction obtained through the matrix factorization $\boldsymbol{WH}$. Through this factorization, linear combinations of the columns of $\boldsymbol{W}$ are produced using weights from $\boldsymbol{H}.$ The Frobenius norm between $\boldsymbol{X}$ and $\boldsymbol{WH}$ was used in our calculations, with the divergence between the two being calculated as $\frac{1}{2}||\boldsymbol{X} - \boldsymbol{WH}||_2$. Thus, in order to estimate the error associated with each embedding dimension, this divergence or reconstruction error was plotted. For all calculations, each channel in the data set was min-max rescaled to a 0 to 1 range to ensure that only positive elements were included in $\boldsymbol{X}$. All calculations were performed using Scikit-learn[26].

PCA. Principal components analysis (PCA) is a linear dimension reduction method that aims to capture the primary axes of variation in the data on a global level[22]. In order to determine the intrinsic dimensionality of the data set estimated by PCA, the cumulative percentage of residual variance remaining after dimension reduction for each component is plotted. Given a component $1 \leq d \leq n - 1$ where $n$ is the number of dimensions of the original data set, the percentage of variance explained by embedding in dimension $d$ is determined by summing the $d$-largest eigenvalues of the covariance matrix of the full data set. For all calculations, each channel in the data set was standardized by removing the mean and scaling to unit variance. Standardization was used to ensure that no feature dominated the objective function of PCA. All calculations were performed using Scikit-learn[26].

**(iii.) Assessing information content relative to H&E tissue morphology**. In order to have an unbiased assessment of image to image information content between embedded data produced from each dimension reduction method and corresponding H&E stained tissue biopsy sections, three channels from MSI data were carefully chosen as representative peaks that highlighted morphological characteristics of the tissue (*m/z* peaks 782.399, 725.373, 566.770 for diabetic foot ulcer, prostate, and tonsil), a hyperspectral image was created, converted to gray scale, and was registered to the corresponding gray scale converted H&E image (**Supplementary Figs. 1a, 2a, 3a**).

To ensure an appropriate alignment between the manually chosen gray scale MSI image of the diabetic foot ulcer and the gray scale H&E image, the mutual information of the registration and the dice score of seven paired ROIs between the two images were assessed across hyper-parameter grids for an initial affine registration and subsequent nonlinear registration (**Supplementary Fig. 1c**). For the prostate and tonsil tissues, we optimized the mutual information alone (**Supplementary Figs. 2c and 3c**). The results across hyper-parameter grids were then analyzed in order to choose the optimal parameters for each step in the registration scheme.

For the affine registrations, the hyper-parameter search resulted in a chosen number of resolutions in the multi-resolution pyramidal hierarchy. For the nonlinear registrations, both the number of resolutions and final uniform grid-spacing for the B-spline controls points were determined by the hyper-parameter grid search. In both registrations, the number of resolutions either improved registration results or left the registration unchanged. However, during the nonlinear registration, finer control point grid-spacing schedules resulted in improved registrations indicated by the mutual information, yet they resulted in regions with unrealistic warping even with the addition of regularization using deformation bending energy penalties[27]. A value of 300 for the final grid-spacing was chosen as a balance between improved registration indicated by the cost function and increased warping.

The resulting deformation field was then applied to the gray scale hyperspectral images created from each dimension reduction algorithm to spatially align them equally with the H&E images of each tissue. Prior to calculating the mutual information between the H&E and embedded MSI images, a nonzero intersection was applied to the pair of images. The nonzero intersection was used to account for any edge effects introduced in the registration by using three manually chosen MSI peaks, which could have adversely affected the registration and mutual information calculations in our analysis if they were not be well-represented at all locations in the images. The mutual information between each registered dimension reduction image (n = 5 per method) was then calculated using a Parzen window-based method in SimpleITK[28] (**Supplementary Figs. 1b, 2b, 3b**).

**(iv.) Assessing algorithm robustness to noise.** Through the assessment of data intrinsic dimensionality, we learned that both high-dimensional imaging modalities (MSI and IMC) follow a manifold structure, where the dimensionality of the data can be approximated with fewer degrees of freedom than the number of parameters initially given in the ambient space. Using this information, in addition to the visual quality of subsequent spatial mappings of each method back onto tissues, as evidence to justify the assumption of such manifold structure, we then proceeded to interrogate the ability of each algorithm to preserve geodesic distances in low dimensional embeddings with and without the addition of "noisy" peaks and/or technical variation.

To do this, we utilized the denoised manifold preservation (DEMaP) metric[19]. By computing the DEMaP metric (Spearman's rank correlation coefficient) between geodesic distances in the ambient space of a peak-picked MSI data set and the pairwise embedded Euclidean distances between data points from the corresponding non-peak-picked data set, we assessed the ability of each algorithm to preserve the manifold structure of the data set in the presence of noise. Since all the algorithms used were either calculated using the Euclidean metric with 15 nearest neighbors or they inherently assume a Euclidean structure, we calculated geodesic distances in the peak picked MSI data set using 15 nearest neighbors using the Euclidean metric. Peak-picking was performed in SCiLS Lab 2018b using orthogonal matching pursuit with a maximum number of peaks of 1,000. The DEMaP scores for each method across 5 random initializations of each algorithm for each MSI data set are shown in **Supplementary Figs. 1g, 2g, and 3g**.

**(v.) Assessing computational runtime.** Computational runtime for all methods was captured across 5 randomly initialized runs for each algorithm for embedding dimensions 1-10 across diabetic foot ulcer, prostate cancer, and tonsil tissue biopsy MSI data (**Supplementary Figs. 1h, 2h, 3h**).

#### Supplementary Notes References


10. Schapiro, D. et al. MCMICRO: A scalable, modular image-processing pipeline for multiplexed tissue imaging. *bioRxiv* (2021).
11. Schapiro, D. et al. histoCAT: analysis of cell phenotypes and interactions in multiplex image cytometry data. *Nature methods* **14**, 873 (2017).
12. Berg, S. et al. ilastik: Interactive machine learning for (bio) image analysis. *Nature Methods*, 1-7 (2019).
13. Schindelin, J. et al. Fiji: an open-source platform for biological-image analysis. *Nature methods* **9**, 676-682 (2012).
14. Bankhead, P. et al. QuPath: Open source software for digital pathology image analysis. *Scientific reports* **7**, 1-7 (2017).
15. Sofroniew, N., Talley Lambert, Evans, K., Nunez-Iglesias, J., Yamauchi, K., Solak, A. C., Buckley, G., Bokota, G., Tung, T., Ziyangczi, Freeman, J., Boone, P., Winston, P., Loic Royer, Har-Gil, H., Axelrod, S., Rokem, A., Bryant, Hector, Mars Huang, Pranathi Vemuri, Dunham, R., Jakirkham, Siqueira, A. D., Bhavya Chopra, Wood, C., Gohlke, C., Bennett, D., DragaDoncila & Perlman, E. napari/napari: 0.3.5. *(Zenodo, 2020).* (2020).
16. Staring, M., Van Der Heide, U.A., Klein, S., Viergever, M.A. & Pluim, J.P. Registration of cervical MRI using multifeature mutual information. *IEEE transactions on medical imaging* **28**, 1412-1421 (2009).
17. Maaten, L.v.d. & Hinton, G. Visualizing data using t-SNE. *Journal of machine learning research* **9**, 2579-2605 (2008).
18. McInnes, L., Healy, J. & Melville, J. Umap: Uniform manifold approximation and projection for dimension reduction. *arXiv preprint arXiv:1802.03426* (2018).
19. Moon, K.R. et al. Visualizing structure and transitions in high-dimensional biological data. *Nature Biotechnology* **37**, 1482-1492 (2019).
20. Tenenbaum, J.B., De Silva, V. & Langford, J.C. A global geometric framework for nonlinear dimensionality reduction. *science* **290**, 2319-2323 (2000).
21. Kim, H. & Park, H. Sparse non-negative matrix factorizations via alternating non-negativity-constrained least squares for microarray data analysis. *Bioinformatics* **23**, 1495-1502 (2007).
22. Jolliffe, I.T. & Cadima, J. Principal component analysis: a review and recent developments. *Philosophical Transactions of the Royal Society A: Mathematical, Physical and Engineering Sciences* **374**, 20150202 (2016).
23. Leland, M., John, H., Nathaniel, S. & Lukas, G. UMAP: Uniform Manifold Approximation and Projection. *Journal of Open Source Software* **3**, 861 (2018).
24. Connor Meehan, S.M., and Wayne Moore (https://www.mathworks.com/matlabcentral/fileexchange/71902, MATLAB Central File Exchange.; 2020).
25. Ulyanov, D. Multicore-tsne. *GitHub Repos. GitHub* (2016).
26. Pedregosa, F. et al. Scikit-learn: Machine learning in Python. *Journal of machine learning research* **12**, 2825-2830 (2011).
27. Rueckert, D. et al. Nonrigid registration using free-form deformations: application to breast MR images. *IEEE transactions on medical imaging* **18**, 712-721 (1999).

28. Lowekamp, B.C., Chen, D.T., Ibáñez, L. & Blezek, D. The design of SimpleITK. *Frontiers in neuroinformatics* **7**, 45 (2013).
